# Heterodimeric insecticidal peptide provides new insights into the molecular and functional diversity of ant venoms

**DOI:** 10.1101/2020.07.29.226878

**Authors:** Axel Touchard, Helen C. Mendel, Isabelle Boulogne, Volker Herzig, Nayara Braga Emidio, Glenn F. King, Mathilde Triquigneaux, Lucie Jaquillard, Rémy Béroud, Michel De Waard, Olivier Delalande, Alain Dejean, Markus Muttenthaler, Christophe Duplais

## Abstract

Ants use venom for predation, defence and communication, however, the molecular diversity, function and potential applications of ant venom remains understudied compared to other venomous lineages such as arachnids, snakes and cone snails. In this work, we used a multidisciplinary approach that encompassed field work, proteomics, sequencing, chemical synthesis, structural analysis, molecular modelling, stability studies, and a series of *in vitro* and *in vivo* bioassays to investigate the molecular diversity of the venom of the Amazonian *Pseudomyrmex penetrator* ants. We isolated a potent insecticidal heterodimeric peptide Δ-pseudomyrmecitoxin-Pp1a (Δ-PSDTX-Pp1a) composed of a 27-residue long A-chain and a 33-residue long B-chain crosslinked by two disulfide bonds in an antiparallel orientation. We chemically synthesised Δ-PSDTX-Pp1a, its corresponding parallel AA and BB homodimers, and its monomeric chains and demonstrated that Δ-PSDTX-Pp1a had the most potent insecticidal effects in blow fly assays (LD_50_ = 3 nM). Molecular modelling and circular dichroism studies revealed strong alpha-helical features, indicating its cytotoxic effects could derive from membrane disruption, which was further supported by insect cell calcium assays. The native heterodimer was also substantially more stable against proteolytic degradation (t_1/2_ =13 h) than its homodimers or monomers (t_1/2_ <20 min), indicating an evolutionary advantage of the more complex structure. The proteomic analysis of *Pseudomyrmex penetrator* venom and in-depth characterisation of Δ-PSDTX-Pp1a provide novel insights in the structural complexity of ant venom, and further exemplifies how nature exploits disulfide-bond formation and dimerization to gain an evolutionary advantage *via* improved stability; a concept that is also highly relevant for the design and development of peptide therapeutics, molecular probes and bioinsecticides.

## Introduction

Hymenopterans are a large order of insects with ~120,000 described species and over 250 million years of evolution [1,2,3]. Many of their members, including ants, bees and wasps, use venom for predation, defence and communication. These venoms seem to be highly heterogeneous and structurally complex, with a wide range of bioactive constituents being reported including sugars, formic acid, biogenic amines, polyamines, alkaloids, and peptides [4,5]. Considering this immense chemical diversity and the high species richness of this order, hymenopterans can be considered a vast, yet understudied resource for the discovery of new biochemicals that complements venom from other, better studied species such as spiders, scorpions, snakes and cone snails. A systematic analysis of the chemical and structural diversity within hymenopteran venoms does not exist [4,5]. However, the high diversity of ant species with diverse ecology and evolutionary history predicts enormous potential for the discovery of bioactive peptides with novel structural scaffolds and pharmacology with applications in medicine and agriculture [6,7]. This potential has recently been illustrated by the discovery of a structurally unique ion channel ligand from the venom of the ant *Anochetus emarginatus* [8].

Ants belonging to the genus *Pseudomyrmex* possess venoms that rapidly subdue prey and effectively deter herbivores, suggesting that they contain both neurotoxic and cytotoxic compounds [9]. They employ their venoms according to their nesting mode (i.e., terrestrial and arboreal species, and, among the latter, plant-ants or obligate inhabitants of myrmecophytes) [10]. A previous mass spectrometry (MS)-based survey of three *Pseudomyrmex* venoms revealed that the plant-ant species *Pseudomyrmex penetrator* (*P. penetrator*) contains uncharacterized linear peptides as well as disulfide-rich peptides, indicating a complex structural diversity of toxins that warrants further investigation [11].

Here, we studied the venom of *P. penetrator* through proteomics, cytotoxicity-guided venom fractionation, chemical synthesis, structure-activity relationship (SAR) studies, proteolytic stability assays, and *in vivo* characterisation of insecticidal activity.

## Results

### Venom collection, mass spectrometry analysis and cytotoxicity-guided peptide isolation

A total of 11.5 mg of dried crude venom was obtained by dissecting the venom sacs from 609 *P. penetrator* workers (19 μg of dried venom *per* individual). The total amount was substantial considering the small body size of *P. penetrator* workers (~5 mm) and is linked to the relatively large size of the venom sac, which is ~0.5 mm and occupies a large volume within the ant gaster (**Fig. 1 A** and **B**).

**Fig. 1.**
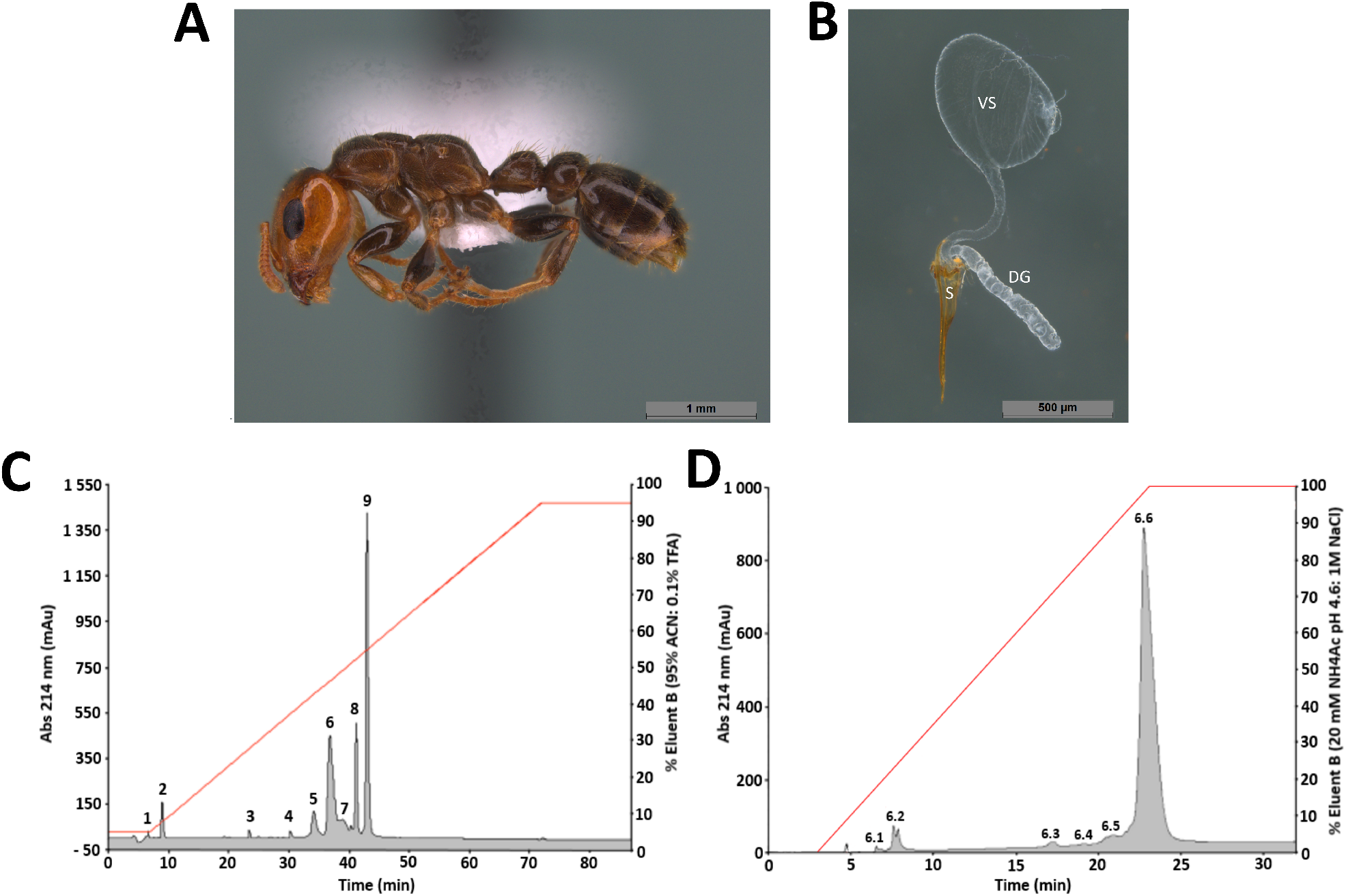
Venom peptide discovery from *P. penetrator* ants. **A** Photo of a *P. penetrator* worker. **B** The venom sac (VS), Dufour’s gland (DG) and sting (S) after dissection. **C** RP-HPLC chromatogram resulting from of *P. penetrator* crude venom fractionation on a C_18_ column. Fractions labelled from 1 to 9 were selected for cytotoxicity assays on insect cells. Fraction 6 exhibited the highest cytotoxic activity. **D** Cation-exchange chromatogram of RP-HPLC fraction 6. Sub-fractions were labelled 6.1 to 6.6. Sub-fraction 6.6 represented > 90% of the total fraction 6 content and contained the pure cytotoxic peptide Δ-PSDTX-Pp1a.

The cytotoxicity of the crude venom was assessed by measuring growth inhibition of *Aedes albopictus* mosquito C6/36 cells. *A. albopictus* is an important vector of mosquito-borne diseases, including dengue, yellow fever, chikungunya and Zika, and its widespread geographic redistribution puts a large population at risk of contracting these diseases [12]. *A. albopictus* C6/36 cells are therefore an important *in vitro* model for the discovery of novel cytotoxic compounds and development of novel insecticidal agents. Crude *P. penetrator* venom had potent cytotoxic activity with an IC_50_ of 2 μg/mL (**Table S1**). The crude venom was then screened in a cytometry assay to examine the effects on intracellular concentrations of calcium (Ca^2+^), sodium (Na^+^) and chloride (Cl^−^) ions in *A. albopictus* C6/36 cultures. Higher intracellular Ca^2+^ concentrations were observed in cells treated with *P. penetrator* crude venom (**Fig. 2**). At very low concentrations (0.03 and 0.3 μg/mL), the venom extract significantly elevated intracellular Ca^2+^ concentration (p-value <0.0001 and <0.05 respectively). No effects were observed for Na^+^ and Cl^−^ ions.

**Fig. 2.**
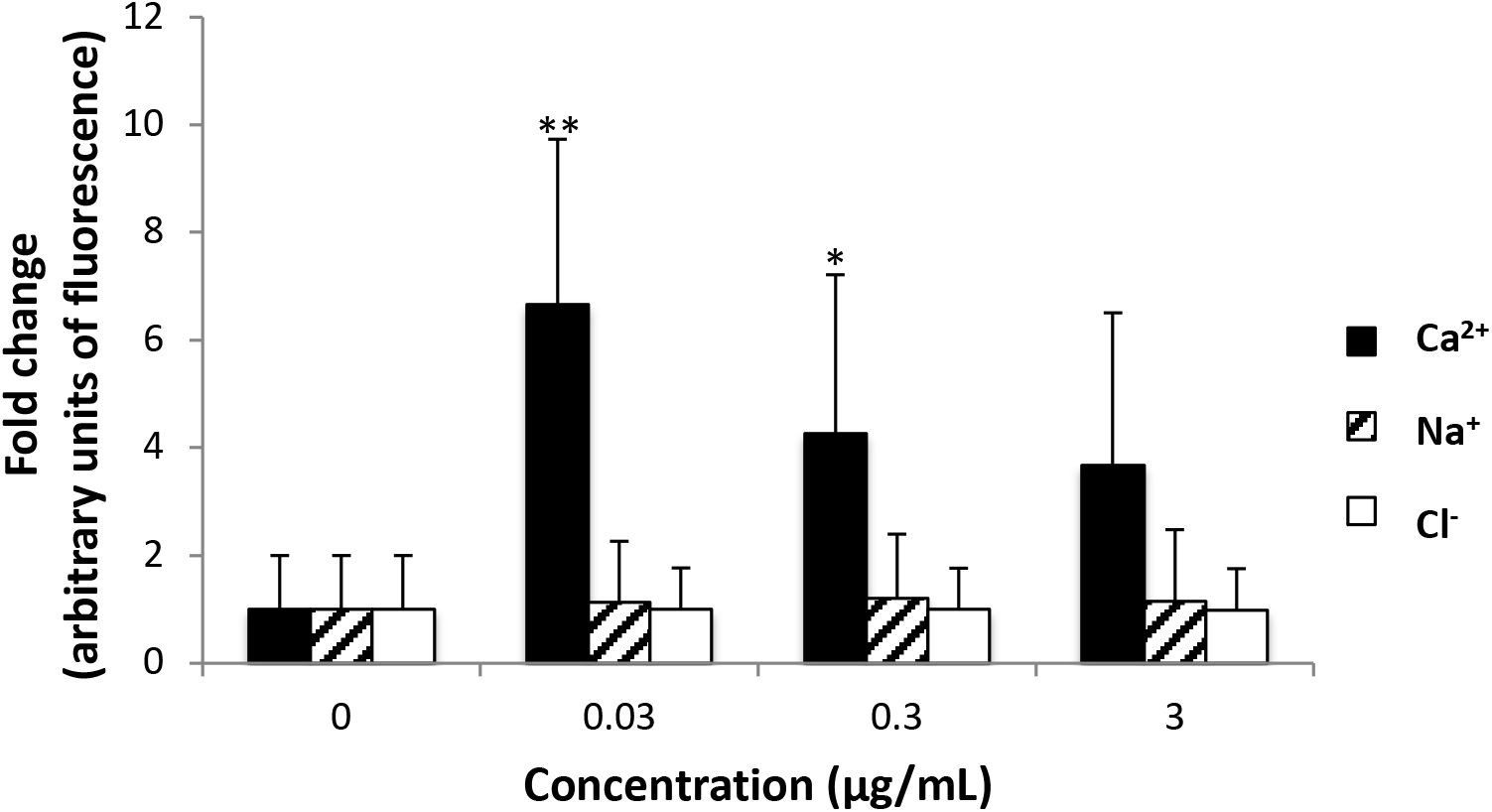
Cytometry assay of crude *P. penetrator* venom. Fold change in fluorescence intensity as a proxy for intracellular Ca^2+^ (black box), Na^+^ (dashed box), Cl^−^ (white box) concentrations in *A. albopictus* C6/36 cells treated with different concentrations of crude *P. penetrator* venom. Asterisks indicate significant differences based on non-parametric analysis using the Kruskal-Wallis test with Dunn’s multiple comparison test (with ** for p-values <0.0001 and * for p-values < 0.05). Results based on four independent experiments (standard deviation error bars).

Fractionation of the crude venom using reversed-phase high-performance liquid chromatography (RP-HPLC) revealed nine peaks, with just three main (peaks 6, 8 and 9) constituting >60% of the crude venom mass (**Fig. 1C**). The monoisotopic masses of peptides in peaks 4–9 were determined by electrospray ionisation quadrupole time-of-flight mass spectroscopy (ESI-Q-TOF MS) and compared with the monoisotopic masses identified in a previous study [11] (**Table S1**). Peaks 1-3 had no defined mass and were not tested in the bioassays. A total of sixteen different peptide masses were identified ranging from 649.4–7242.1 Da including three homo- and heterodimeric peptides (5955.4, 6598.8, 7242.1 Da). Nine of the sixteen masses were previously reported [11] (**Table S1**).

The cytotoxicity of the nine RP-HPLC fractions were tested on *A. albopictus* cells (**Table S2**). Five of these fractions were cytotoxic (fractions 5 to 9). Fraction 6 was the most potent one with cytotoxic activity (IC_50_ 3.16 μg/mL) similar to the whole crude venom. Fraction 6 was further sub-fractionated by cation exchange chromatography and a heterodimeric peptide with a monoisotopic mass of 6598.8 Da was isolated (**Fig. 1D**, **Fig. S2**). This newly discovered ant venom peptide was highly cytotoxic to *A. albopictus* cells (IC_50_ 1.04 μM) and named Δ-pseudomyrmecitoxin-Pp1a (Δ-PSDTX-Pp1a) following established nomenclature [4]. Δ indicates peptides with cytolytic activity, PSDTX denotes peptides from ants of the subfamily Pseudomyrmecinae, ‘Pp’ are genus/species descriptors, and “1a” is for distinguishing paralogous peptides from the same venom [13].

### Δ-PSDTX-Pp1a sequence determination, chemical synthesis and *in silico* modelling

Δ-PSDTX-Pp1a was sequenced using a combination of chemical reduction with dithiothreitol, Edman degradation, enzymatic digestion and *de novo* MS sequencing (**Table S3**). This yielded the sequence of a heterodimeric peptide comprised of a 27-residue A-chain with a C-terminal amide and a 33-residue B-chain with a C-terminal acid. Each chain contains two cysteine residues, which covalently link the two chains through two disulfide bonds. Enzymatic degradation and MS analysis revealed an antiparallel orientation of the heterodimer with a disulfide bond connectivity of Cys_A_^I^-Cys_B_^II^ and Cys_A_^II^-Cys_B_^I^ (**Fig. 3**). Both peptide chains are highly cationic (predicted isoelectric point of 9.93 for A-chain and 9.70 for B-chain) and highly homologous to each other (85% sequence identity). In addition to being highly cationic (net charge of +7 for the A-chain and +6 for the B-chain) they are amphiphilic (41% and 45% hydrophobic residues, for A- and B-chain, respectively) (**Fig. S1**).

**Fig. 3.**
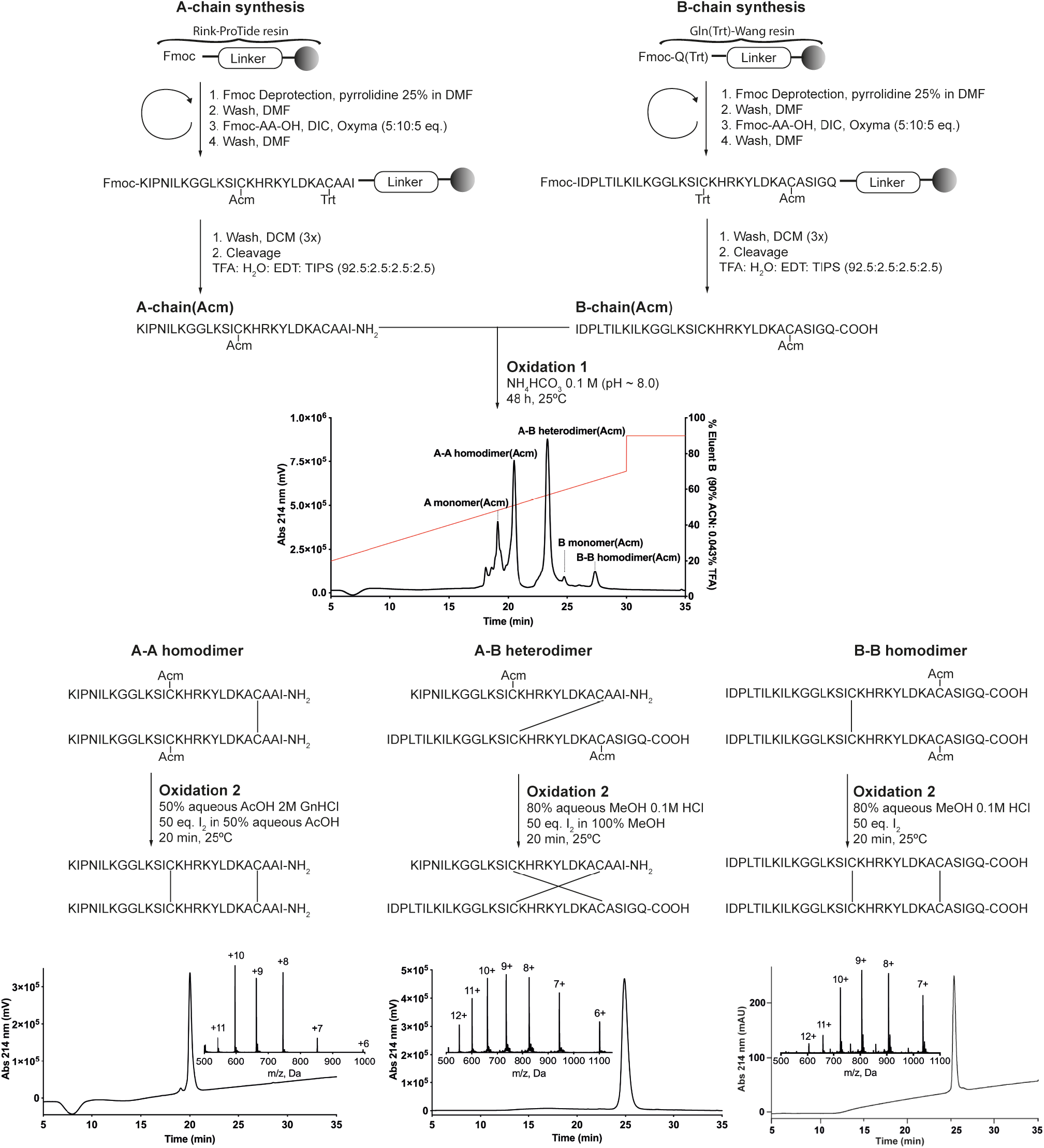
Chemical synthesis of antiparallel heterodimeric Δ-PSDTX-Pp1a and the A-chain and B-chain homodimers. A-chain and B-chain peptides were assembled using Fmoc-SPPS and purified using RP-HPLC. Cysteine residues were differentially protected: A-chain Cys_A_^I^ and B-chain Cys_B_^II^ were protected with acetamidomethyl (Acm) and A-chain Cys_A_^II^ and B-chain Cys_B_^I^ with trityl (Trt). Upon treatment with trifluoroacetic acid (TFA), the peptides were deprotected and released from the resin; Cys_A_^I^ and Cys_B_^II^ remained protected with Acm. Subsequent folding (oxidation 1) produced the single disulfide bond connected A-B heterodimer and the A-A and B-B homodimers. Acm deprotection and formation of the second interchain disulfide bond (oxidation 2) produced the fully folded A-B heterodimer and A-A and B-B parallel homodimers. After cleavage and after each oxidation, peptides were purified *via* preparative C_18_ RP-HPLC, lyophilised and analysed using analytical RP-HPLC and MS. Measured isotope peaks for the final dimer products are listed in Table S4.

To further study Δ-PSDTX-Pp1a, we chemically synthesized this peptide *via* Fmoc-SPPS (9-fluorenymethyloxycarbonyl-solid phase peptide synthesis) in combination with a directed folding strategy using orthogonally protected cysteine building blocks with acetamidomethyl (Acm) and trityl (Trt) groups (**Fig. 3**). The first disulfide bond formation and dimerization step was carried out in a 1:1.5 ratio mixture of reduced A- and B-chains in aqueous buffer (0.5:0.75 mM A-chain:B-chain, 0.1 M NH_4_HCO_3_, 25°C, pH 8.3, 48 h) resulting in the expected three products, the A-B heterodimer and the two homodimers (A-A, B-B), all connected by a single bond. Following RP-HPLC purification, each product was subjected to iodine oxidation to remove the Acm group and to form the second interchain disulfide bond, thereby producing the fully folded antiparallel heterodimer and the two parallel homodimers. The synthetic heterodimer was confirmed to be identical to native Δ-PSDTX-Pp1a by a RP-HPLC coelution study and comparison of the high-resolution mass and MS fragmentation patterns (**Fig. S2**). In addition to the dimeric peptides, the single A- and B-chain were synthesized using the same Fmoc-SPPS methodology but without Acm protected cysteine residues to obtain the reduced linear A- and B-chain monomers (**Fig. S3, S4**).

To assess whether the structural or surface properties of the hetero- and homodimer peptides could affect their activity and stability, Δ-PSDTX-Pp1a and the parallel A- and B-chain homodimers were modelled *in silico* using the *de novo* PEP-fold method [14] and molecular dynamics (MD) using the Amber forcefield as implemented in Yasara [15] (**Fig. 4**). The A-chain was predicted to have a N-terminal 3_10_-helix (Iso_A_2-Lys_A_7) and an α-helix (Leu_A_10-Lys_A_22) while the B-chain was predicted to contain two successive α-helices (Pro_B_3-Gly_B_12 and Leu_B_14-Iso_B_31) (**Fig. 4A**). The C-terminal α-helices of Δ-PSDTX-Pp1a are packed together in a compact non-coiled coil structure stabilized by two disulfide bonds (**Fig. 4B**). Most of the hydrophobic and hydrophilic residues are largely surface exposed even though the lateral chain of Asp_B_25 is buried in the cationic core of the A-B heterodimer (**Fig. 4C**). This negatively charged residue likely forms a strong electrostatic interaction with the Lys_A_15 and Lys_A_18 of the A chain and contributes to inter-chain stability and facilitates close packing of the C-terminal α-helices. The Δ-PSDTX-Pp1a heterodimer is highly cationic with clusters of solvent-exposed Lys, Arg, His and Ser side chains, leading to an overall hydrophilic surface on one side of the heterodimer. By contrast, the other side is mostly hydrophobic except for a central hydrophilic core (**Fig. 4D**). A circular dichroism (CD) study of Δ-PSDTX-Pp1a confirmed the presence of the predicted α-helical secondary structure *via* a positive band at 190 nm and two negative bands at 208 nm and 222 nm (**Fig. 4E**).

**Fig. 4.**
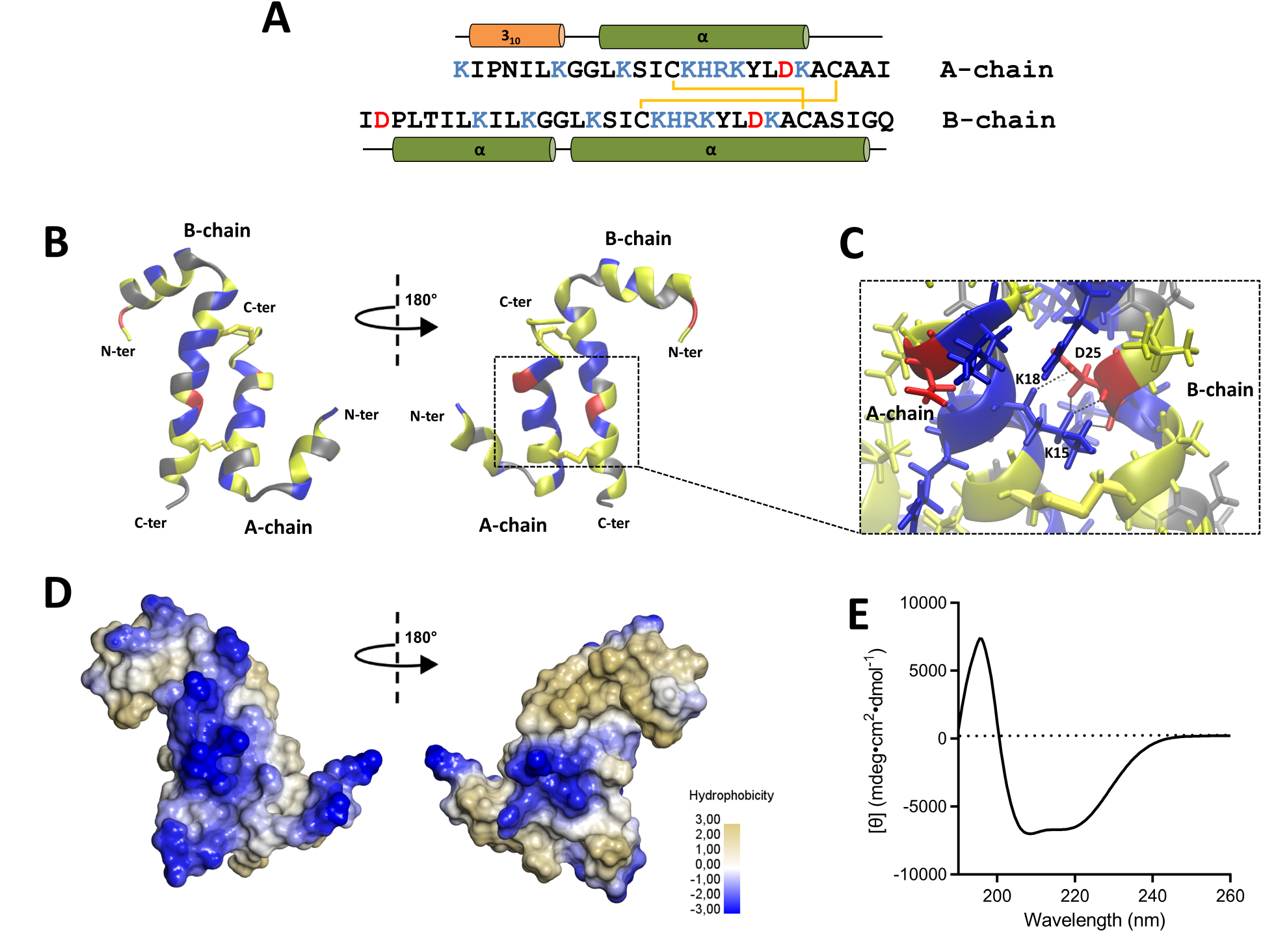
Structural analysis of Δ-PSDTX-Pp1a. **A** Amino acid sequence of Δ-PSDTX-Pp1a with cationic and anionic residues in blue and red, respectively. Schematic representation of the predicted secondary structure of Δ-PSDTX-Pp1a is shown above and below the sequences. **B** Most representative geometry for the heterodimeric Δ-PSDTX-Pp1a peptide after clustering of a 40 ns molecular dynamics simulation. The cationic residues are in blue, anionic residues in red, polar non-charged residues in grey and hydrophobic amino acids in yellow. **C** Expanded view of the central hydrophilic core showing the buried side chain of Asp_B_25. **D** Molecular surface representation of Δ-PSDTX-Pp1a highlighting the predominance of hydrophilic patches. **E** CD spectra of Δ-PSDTX-Pp1a dissolved in sodium phosphate, pH 7.4.

Notably, the predicted structure of the two homodimers revealed a disordered helical conformation and a decrease of overall α-helicity in comparison to the antiparallel heterodimer (**Fig. S5** and **S6**). These disordered helical structures of both homodimers appear to be the consequence of negative interactions between similar charged residues (**Fig. S5A-D**). The BB-homodimer has similar surface properties to Δ-PSDTX-Pp1a, albeit the hydrophobic patches are more dispersed, while the AA-homodimer has the least hydrophobic surface. Based on this structural characterization, the three dimers appear to have an amphipathic character where cationic residues represent a discontinuity within the hydrophobic patches. Therefore, it was hypothesized that the cytotoxicity activity of Δ-PSDTX-Pp1a arises from an interaction with cell membranes as often reported with such cationic amphipathic α-helical peptides [16,17].

### Insecticidal and stability study

To gain insights into the ecological role and potential evolutionary advantage of these heterodimeric peptides, a structural class of toxins commonly reported in ant venoms, we embarked on a comparative investigation of insecticidal activity and stability of the Δ-PSDTX-Pp1a heterodimer along with the two parallel homodimers and the A- and B-chain reduced monomers. Insecticidal activity was tested *in vivo* by intrathoracic injection into sheep blowflies (*Lucilia cuprina*), a serious agricultural pest commonly used as a model organism to monitor the insecticidal activity of venom peptides [18]. The Δ-PSDTX-Pp1a heterodimer as well as the A- and B-chain homodimers exhibited the most potent insecticidal effects, with both A- and B-chain monomers being significantly less potent (**Fig. 5** and **Table S5**). For the homo- and heterodimeric peptides, contractile paralysis developed almost instantly while the flies were still attached to the injection needle and was fully developed when fly behaviour was first measured at 30 min after injection. At higher doses, paralysed flies did not recover with most being dead at 24 h post-injection.

**Fig. 5.**
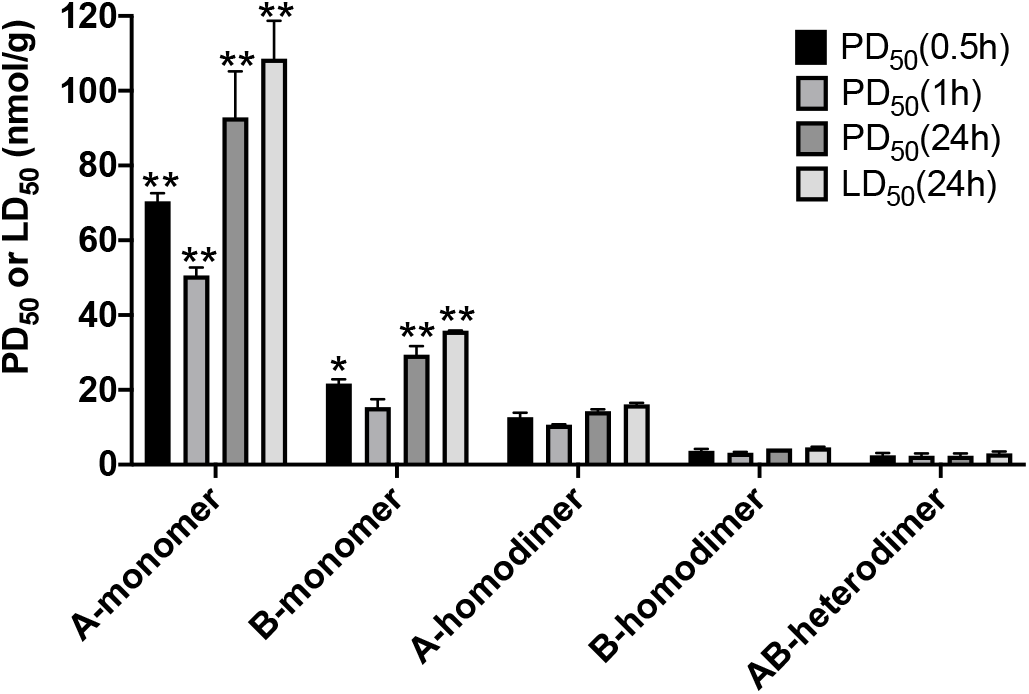
Insecticidal activity of Δ-PSDTX-Pp1a. Paralytic and lethal effects of monomeric (A, B), homodimeric (AA, BB) and heterodimeric (AB) Δ-PSDTX-Pp1a analogues when injected into sheep blowflies (*L. cuprina*). Statistical significance is based on a two-way ANOVA followed by Tukey’s post hoc test and indicated by * (p<0.01) and ** (p<0.0001) as compared against the respective PD_50_ and LD_50_ values of the heterodimeric Δ-PSDTX-Pp1a. Error bars represent the standard error of the mean.

We then tested the proteolytic stability of the peptides by incubating them with proteinase K, a broad-spectrum serine protease which cleaves peptides bonds at the C-terminal side of aromatic and aliphatic residues and is used to test protein/peptide stability as it typically (37°C, pH = 7.5). Both the monomers and parallel homodimers were degraded within 20 min with nearly identical kinetics. By contrast, the heterodimer was exceptionally resistant to proteolytic degradation with a half-life (t_1/2_) of 13 h, making it >39-fold more stable than the monomers and homodimers. This is particularly impressive considering that the heterodimer contains 24 potential proteinase K cleavage sites dispersed throughout both peptide chains (**Fig. S7**). Δ-PSDTX-Pp1a was also very stable to heat, with a stable half-life of ~13 h at 90°C (**Fig. S8**).

## Discussion

Ant species that inhabit plants (myrmecophytes) in an obligatory mutualism use a defensive venom to protect the host against defoliating insects and browsing mammals. Consequently, their venom has evolved toxins that trigger pain in vertebrates and paralyse/kill large arthropods (e.g., caterpillars, grasshoppers). Previous investigations conducted on two other plant-ant venoms (i.e., *P. triplarinus* and *Tetraponera aethiops*) revealed the presence of uncharacterized disulfide-linked dimeric peptides, suggesting that this class of toxin may have been retained in *Pseudomyrmecinae* because it participates in host plant protection [19,20]. Dimeric peptides have rarely been reported in venom peptidomes, with only a few examples in snake [21,22], scorpion [23], spider [24] and cone snail [25] venoms. Despite the fact that only a few studies have examined ant venom peptidomes, a total of 22 homo- and heterodimeric peptides have been identified from the venoms of the ant subfamilies Ectatomminae [26], Myrmeciinae [17,27], Pseudomyrmecinae [19,20] and Ponerinae [28], indicating that these structurally more complex peptides play an important role in ant venom. From an evolutionary perspective, the presence of dimeric peptides within the venom of these four ant subfamilies is also highly interesting since they constitute a non-monophyletic group [29], leading to the question on what are the evolutionary advantages of these dimeric toxins compared to monomeric toxins?

We thus set out to study the venom of *P. penetrator*, a plant-ant species that strictly uses its venom to defend the host tree [10] and revealed that the most cytotoxic component of the whole venom was Δ-PSDTX-Pp1a, a novel antiparallel heterodimer peptide. We then chemically synthesized Δ-PSDTX-Pp1a along with its homodimeric and monomeric analogues to provide an in-depth characterization of this toxin in terms of function, structure, stability and potential applications.

### Insecticidal activity

Δ-PSDTX-Pp1a is a fast-acting insecticidal peptide which caused immediate paralysis when injected into blowflies leading to death within 24 h. Compared to monomeric venom peptides of other ant species, Δ-PSDTX-Pp1a with a LD_50_ (at 24 h) of 3.0 nmol/g was more lethal than the most potent insecticidal venom peptide from *Manica rubida* (LD_50_ at 24 h = 75.45 nmol/g for U_20_-MYRTX-Mri1a, tested in *Lucilia caesar*) [30] and from *Neoponera goeldii* (LD_50_ at 24 h = 25.7 nmol/g for ponericin G1, tested in *Acheta domesticus*) [31]. To date, most of the insecticidal bioassays conducted on ant venoms revealed that the lethality of peptides is relatively weak since most exhibit non-lethal paralytic effects which are often reversible [8, 17, 30]. For instance, PONTX-Ae1a toxin isolated from the venom of the predatory ant *Anochetus emarginatus* [8] displayed similar paralytic activity to Δ-PSDTX-Pp1a on blowflies (PD_50_ at 1 h = 2.4 nmol/g for Δ-PSDTX-Pp1a and PD_50_ at 1 h = 8.9 nmol/g for PONTX-Ae1a), but this activity was completely reversible at all doses. The high lethality observed for Δ-PSDTX-Pp1a could be advantageous regarding the non-predatory behaviour of *P. penetrator* that uses its venom for long-term protection of the host-plant while predatory ants store living paralysed prey in their nest before consumption [32,33]. In a broader context, the insecticidal potency of Δ-PSDTX-Pp1a is about an order of magnitude less potent than the most spider-venom peptides that have been tested in the same blowfly toxicity assay [34,35,36].

Our structure-function relationship studies support a functional gain in lethality with dimerization, in comparison to the monomeric A- and B-chains that are only weakly active, which aligns well with a recent study on heterodimeric ant venom peptide Mp1a from *Ectatomma tuberculatum* ant [37] and a study on homotarsinin, a homodimeric peptide isolated from skin secretions of *Phyllomedusa tarsius* [38].

### Stability

Δ-PSDTX-Pp1a has exceptional proteolytic and thermal stability **(Fig 6, S8)**. The proteolytic stability of the heterodimer (t_1/2_ = 13 h) depends directly on its evolutionarily derived secondary structure considering that the sequence holds 24 potential proteinase K cleavage sites and that the parallel homodimers were degraded 39-fold more quickly (t_1/2_ < 20 min). The interplay between α-helical conformation, disulfide bonds and interfacial hydrophobic interactions of dimeric peptides can be crucial components for high proteolytic stability [42]. Distinctin, for example, a heterodimeric pore-forming peptide found in skin secretions of the frog *Phyllomedusa distincta,* is also resistant to proteolysis [39, 40]. In water, distinctin forms a non-covalent four-α-helix bundle having a positively charged surface and a hydrophobic core [41]. Such non-covalent structural arrangements could also be at play with Δ-PSDTX-Pp1a.

**Fig. 6.**
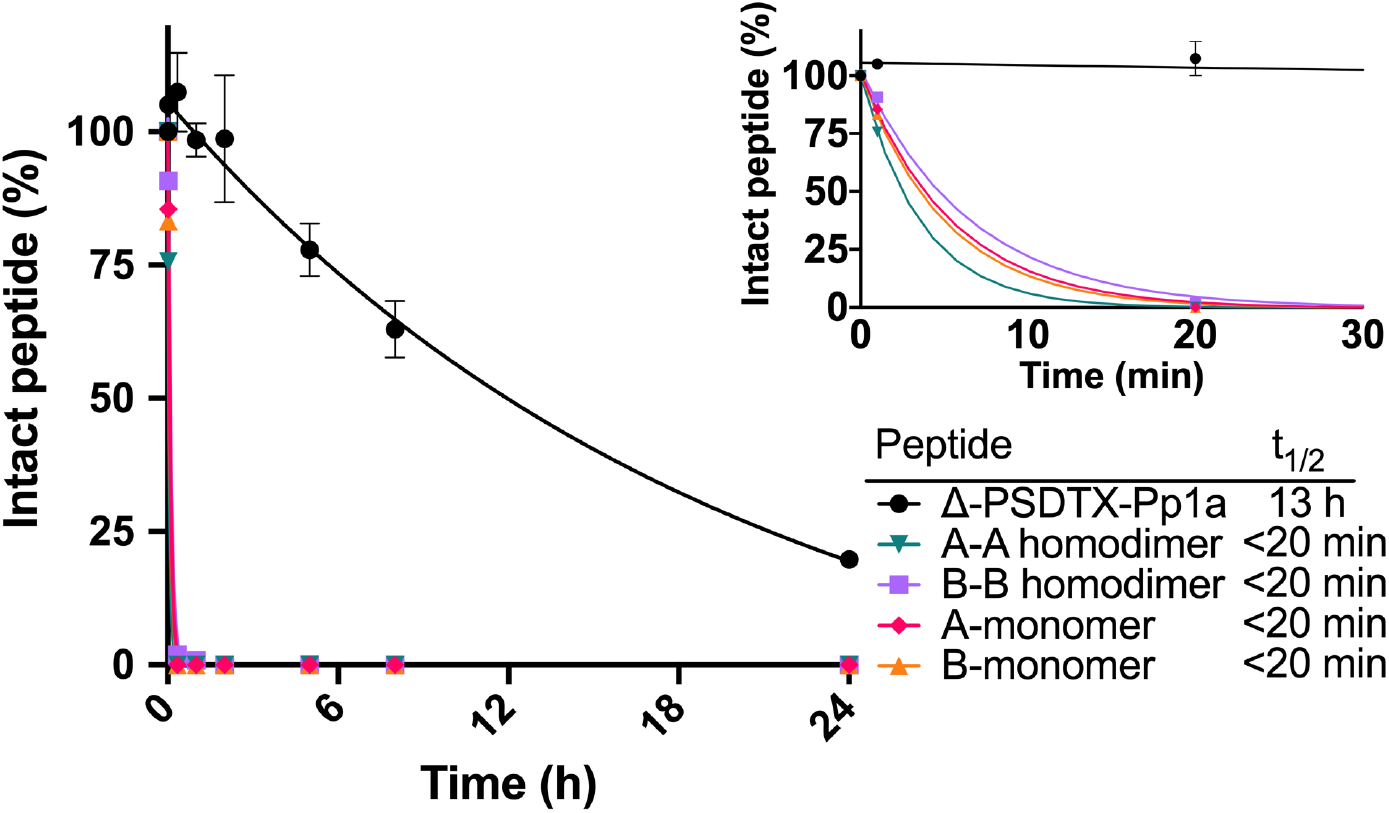
Proteolytic stability of synthetic Δ-PSDTX-Pp1a, AA-homodimer, BB-homodimer, A-monomer and B-monomer. Fraction of intact peptide after incubation with proteinase K (1:200 proteinase K:peptide molar ratio) at pH 7.5 and 37°C for up to 24 h (inset shows the first 30 min). As a negative control, peptides were incubated for 24 h under the same conditions without proteinase K. Peptide values were quantified relative to the negative control at t=0h. N = 2 for synthetic Δ-PSDTX-Pp1a for A-monomer, B-monomer, and AA-homodimer, N = 1 for BB-homodimer. Experiments were performed in duplicate. Data analysis and half-life (t_1/2_) calculation was performed using a non-linear fit one-phase decay model using Prism Version 8. Note the t=24 h error bars of Δ-PSDTX-Pp1a are smaller than the symbol.

There are several other structural features that have evolved in toxins to convey metabolic stability, which is important in order to effectively reach the site of action, which is often the central nervous system of animals. The inhibitor cystine knot (ICK) fold, for example, is widely found in venom peptides [43] and imparts peptides with remarkable stability against proteases [44]. Another robust peptide fold that imparts high protease and thermal stability is the helical arthropod-neuropeptide-derived (HAND) scaffold found in some spider and centipede venoms [45]. It is now well established that several ants use linear and polycationic monomeric peptides to paralyze their prey, and that these peptides are generally inherently unstable and susceptible to proteolytic degradation [46,47,48]. Enzymatic stability is particularly important for venom peptides that have to be injected into a prey/predator and find their *in vivo* target; thus, proteolytic stability could have been a natural selection criterion that promoted the evolution of highly stable dimeric membrane-active peptides in ant venoms.

### Homology, function and molecular target of Δ-PSDTX-Pp1a

The A and B chains of Δ-PSDTX-Pp1a have considerable sequence identity, differing only in their N-terminal region (extension in B chain) or C-terminal region (extensions of 2 and 4 residues) (**Fig. 7**). This high homology suggests that they are encoded by duplicated genes. Compared to other dimeric venom peptides from ants, Δ-PSDTX-Pp1a chains have some sequence homology with the chains of six pseudomyrmecitoxins isolated from the venom of the plant-ant species *P. triplarinus* (31–36% identity and 55–59% similarity) (**Fig. 7**).

**Fig. 7.**
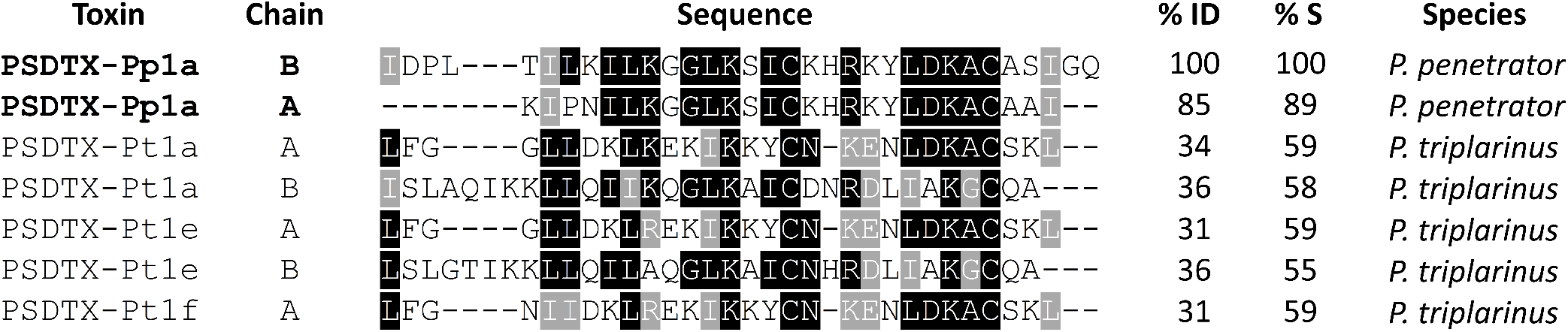
Multiple sequence alignment of subunit chains belonging to dimeric pseudomyrmecitoxins. Gaps were introduced to optimise the alignment. Resulting alignments using the T-coffee alignment program were edited with BOXSHADE 3.3.1-9. Identical residues are highlighted in black while similar residues are highlighted in grey. Both percentage identity (% ID) and percentage similarity (% S) are relative to Δ-PSDTX-Pp1a chain B sequenced in this study.

These pseudomyrmecitoxins are heterodimeric polypeptides associated with the anti-inflammatory activity observed in *P. triplarinus* venom [20]. There is no significant sequence homology between Δ-PSDTX-Pp1a and the pseudomyrmecitoxins described in *T. aethiops* as for the other dimeric peptides from ant venoms. However, the predicted 3D structure of Δ-PSDTX-Pp1a aligns well with the 3D structures of two dimeric ant venom peptides, ectatotoxin Et1a (formerly ectatomin) from *Ectatomma tuberculatum* and myrmeciitoxin Mp1a (formerly pilosulin 2) from *Myrmecia pilosula* [17,49]. Indeed, despite very distinctive amino acid sequences compared to Δ-PSDTX-Pp1a, the structures of these toxins are dominated by α-helices stabilized by two or three disulfide bonds [17,49]. All of the dimeric peptides described in ant venoms share several physicochemical properties including a net positive charge (+4 to +18) due to a high lysine content, a substantial proportion of hydrophobic residues (>40%) and a mass of 5–9 kDa. These features are also shared with membrane-active peptides from ant venom that display insecticidal, cytotoxic and antimicrobial activities. Cell membranes are the common molecular target of linear, polycationic and amphiphilic ant-venom peptides [17,31,46,50]. Among ant venoms, the biological activity and the molecular target have been described for very few dimeric peptides. Nevertheless, Et1a and Mp1a, are pore-forming peptides that induce the formation of nonselective cationic channels in cell membranes, increasing cell permeability with resultant ion leakage and finally cell death [17, 51]. Similar interactions with lipid bilayers were also observed with the homodimeric MIITX_1_-Mg2a peptide isolated from the venom of *Myrmecia gulosa*, which produces pain in vertebrates *via* the formation of pores in the membranes of peripheral sensory neurons [17]. Thus, taken together, the data suggest that the cytotoxic activity of Δ-PSDTX-Pp1a is due membrane pore formation.

In conclusion, Δ-Pseudomyrmecitoxin-Pp1a from the venom of *P. penetrator* is a potent cytotoxic and insecticidal heterodimeric peptide that has higher potency and proteolytic stability than its homodimeric and monomeric counterparts, suggesting an evolutionary advantage. This study further supports that venoms of ants (Formicidae) are a promising but underexplored source of chemically diverse bioactive peptides. Particularly, the presence of such structurally complex and highly stable heterodimers, which seems to be more common in ant venoms than in other animal venoms, highlights ant venom as an attractive biosource for interesting new ligands with applications as bioinsecticides or therapeutic leads, where stability towards abiotic (pH, light, water content) and biotic (enzymes, microorganisms) conditions are desired.

## Materials and Methods

### Venoms collection

Live *P. penetrator* workers (N = 600) were collected on La Montagne des Singes (5°04’20″N; 52°41’43″W) in French Guiana. We used pruning scissors to cut up *Tachigali* aff. *paniculata* compound leaves containing parts of *P. penetrator* colonies and placed them in plastic bags. The boxes and plastic bags containing the colonies were immediately transported to the laboratory where workers were separated and sacrificed by freezing. Ant venom reservoirs were dissected and pooled in 10% v/v acetonitrile (ACN)/ distilled water (v/v) (see protocol in [11]). Briefly, samples were centrifugated for 5 min at 12,000 *g*, then the supernatant was collected and lyophilized prior to storage at −20°C. A total of 608 dissected venom sacs were used for venom fractionation, isolation and sequencing of Δ-PSDTX-Pp1a as well as for the cytotoxicity assays.

### Cytotoxic bioassays

*Aedes albopictus* cells C6/36 were kindly provided by the Virology Unit of the Pasteur Institute of French Guiana. Each well of a 96-well plate was filled with 100 μL of insect-cell suspension (age 1 week, concentration 10^5^–10^7^ cells/mL) and the plate was incubated for 24 h at 28°C. After incubation, the supernatant was removed and replaced with 50 μL of L15 Leibovitz culture media (negative control), cypermethrin (a synthetic pyrethroid) at 300, 100 and 50 μg/mL (positive controls), or venom fractions. These fractions were prepared with lyophilized crude venom to obtain final concentrations of 50 to 0.0005 μg/mL in L15 Leibovitz culture media. Plates were incubated then for another 24 h at 28°C. After incubation, 5 μL of the tetrazolium dye 3-(4,5-dimethylthiazol-2-yl)-2,5-diphenyltetrazolium bromide (MTT, 5 mg/mL) was added to each well and the plate was incubated for 1 h at 28°C in darkness. The supernatants were removed and 50 μL of DMSO was added to each well to suspend the formazan crystals that had formed in the cells. After homogenisation by pipetting, absorbance was measured at 570 nm. Cytotoxic effects were determined by comparing the percentage of living cells treated with the extract with the percentage of living cells treated only with the L15 Leibovitz culture media without venom fractions or cypermethrin. The following formula was used: mortality rate = absorbance of the negative control – (absorbance of the sample/absorbance of the negative control) ×100. Inhibition concentrations (IC_50_ and IC_99_) and their 95% confidence intervals were calculated for six technical replicates (in three independent experiments) per concentration with logistic regression *via* probit analysis [52–55].

### Quantification of intracellular calcium, sodium and chloride ions

Intracellular Na^+^, Ca^2+^ and Cl^−^ concentrations were determined using the cell-permeant ion-specific fluorescent dyes Corona Green Sodium Indicator, Oregon Green 488 BAPTA-1 and MQAE, respectively. All assays were performed using fluorescence-activated cell sorting (FACS) and analyzed with Cell Quest Pro software. Each concentration was tested using six technical replicates (in four independent experiments). Each well of a 24-well plate was filled with 720 μL of *A. albopictus* cells C6/36 suspension (age 1 week, concentration 10^5^–10^7^ cells/mL) and the plate was incubated for 24 h at 28°C. After incubation, the supernatant was removed and 1 mL of the crude venom, solubilized in L15 Leibovitz culture media, was added to obtain final concentrations of 0.03, 0.3 and 3 μg/mL. L15 Leibovitz culture media without venom was used as negative control. After 24 h at 28°C, the supernatant was removed. For Ca^2+^, 250 μL of PBS with 63 μL of Oregon green at 40 μM was added. The plate was then incubated for 60 min at 25°C in the dark. For Cl^−^, 500 μL of hypotonic MQAE solution at 5 mM was added. The plate was then incubated for 15 min at 37°C in the dark. For Na^+^, 500 μL of Corona Green at 10 μM was added. The plate was then incubated for 45 min at 28°C in the dark. For all treatments, cells were washed to remove excess probes and then resuspended in 2 mL of PBS before FACS reading according to the manufacturer’s instructions (Life Technologies) [56–59]. Non-parametric analyses were performed for six technical replicates (four independent experiments) per concentration using the Kruskal-Wallis test and multiple comparisons were performed with the Dunn method [49,53].

### RP-HPLC fractionation and peptide purification

*P. penetrator* venom (11 mg) was fractionated *via* RP-HPLC using a semi-preparative-C_18_ Jupiter Proteo column (4μm, 10 x 250 mm) with a gradient comprised of solvent A [water/0.1 % (v/v) TFA] and solvent B [ACN /0.1% (v/v) TFA]. The gradient of solvent B was as follows: 5% for 7 min, 5–95% over 65 min, and 95% over 8 min at a flow rate of 3.5 mL/min. The eluate was monitored by UV absorbance at 214 nm using a diode-array detector. All analyses were performed on a Shimadzu LC-20AD system. 64 fractions of 3.5 mL were collected automatically every 1.4% of the gradient (every minute), lyophilized and stored at −20°C. Further purification of the most cytotoxic fractions was achieved by subjecting the C_18_ fractions to a second purification step using cation exchange chromatography on a TOSOH Bioscience column (TSK gel SP-STAT, 7 μm, 4.6 mm ID x 10 cm L, TOSOH Bioscience, Germany) with solvent A [200 mM sodium acetate, pH 4.6] and solvent B [200 mM sodium acetate, pH 4.6, 1 M sodium chloride]. The gradient of solvent B was as follows: 0% for 3 min, 0–100% over 20 min, and 100% over 8 min at a flow rate of 1 mL/min. The eluate was monitored by UV absorbance at 214 nm using a diode-array detector. All analyses were performed on a Shimadzu LC-20AD system. Fractions collection was based on time, every 0.5 min between 0.5 and 29.5 min (starting at 0% for 2.5 min and then the gradient itself). Major peaks were desalted by RP-HPLC using an Ascentis C_18_ column (3 μm, 4.6mm ID x 15 cm, Sigma-Merck, Germany) using peak-based collection (slope). Sub-fractions were lyophilized for further cytotoxicity assays. The content of each HPLC fraction was analysed using MS as described in **Table S1**.

### Purification and characterization of the heterodimer and sequencing

Reduction of Δ-PSDTX-Pp1a heterodimer was performed by treating 65 μg of the compound in 100 mM (50 μL) of ammonium bicarbonate buffer pH 8.5 with 40 mM tris(2-carboxyethyl)phosphine (TCEP) for 1 h at 55 ºC (volume of 50 μL), and reaction progress was monitored by MS. Monomers were then alkylated with 70 mM iodoacetamide for 1 h in the dark at 25 ºC before adding 240 mM dithiothreitol (final concentration). Finally, the sample was purified *via* RP-HPLC using an Agilent AdvanceBio peptide map column (2.1 × 250 mm) at a flow rate of 0.4 mL/min with solvent A [water/0.1% TFA (v/v)] and solvent B [ACN/0.1% TFA (v/v)]. The gradient of solvent B was as follows: 5% for 3 min, 5–15% over 1 min, 15–65% over 25 min, 95% over 4 min. The monomers were collected on Agilent 1260 HPLC (Agilent Technologies) using peak-based collection (by slope), then lyophilized. Carbamidomethyl derivates of the monomers were digested in 100 mM of ammonium bicarbonate buffer pH 8.5 with chymotrypsin or Lys-C, at a protein:enzyme ratio of 10:1, for 8 h at 37 ºC and further submitted to MS analysis.

### Determination of disulfide bond connectivity

The disulfide framework of the heterodimer was determined by digesting the compound with trypsin (in 100 mM of ammonium bicarbonate buffer pH 8.5 with trypsin at a protein:enzyme ratio of 10:1) in the presence of the oxidizing reagent cystamine to narrow scrambling effects usually observed in alkaline conditions, and therefore to maintain the native configuration.

The antiparallel form, C14(PS1)-C28(PS2) and C24(PS1)-C18(PS2), was detected by MS through the detection of tryptic fragments A and B with masses of 893.4 Da and 1095.5 Da, respectively (data not shown). In contrast, the parallel configuration, C14(PS1)-C18(PS2) and C24(PS1)-C28(PS2), could be inferred from the observation of tryptic fragments C and D, 896.4 Da and 1092.5 Da, respectively.

Experimentally, Cys-containing peptides A and B were observed as predominant signals suggesting that the heterodimer was mainly in the antiparallel form with a connectivity of C14(PS1)-C28(PS2) and C24(PS1)-C18(PS2). However, the detection of fragment C by LC-MS, while minor, indicates that the parallel form may also be present. The further addition of cystamine results in an increase of the fragments A and B from the anti-parallel form in MS analyses, indicating that the parallel form could possibly result from an artefactual recombination of the disulfide bridges (data not shown). However, this cannot be concluded definitively.

### Mass spectrometry

A Waters Q-TOF Xevo G2S mass spectrometer equipped with an Acquity UHPLC system and Lockspray source was used for acquisition of LC-ESI/MS and LC-ESI/MS/MS data. The dimer analysis was made by injection of 100 pmol onto the column using a 1.7 μm Acquity UPLC BEH (2.1 × 150 mm) column (Waters) at a flow rate of 0.8 mL/min with solvent A [water/0.1% formic acid (FA) (v/v)] and solvent B ACN/0.1% FA (v/v)]. The dimer was eluted using the following gradient of solvent B: 5–10% over 0.2 min, 10–70 % over 1.3 min, 70–90% over 0.1 min. Separation of monomers digests was performed using a 1.7 μm Acquity UPLC BEH300 column (Waters, 2.1 × 50 mm) at a flow rate of 0.4 mL/min with solvent A [water/0.1% FA (v/v)] and solvent B [ACN/0.1% FA (v/v)]. Injections of 150 mol of the monomer digests were made onto the column. Peptides were eluted using the following gradient of solvent B: 2% over 12 min, 2–10% over 1.2 min, 10–70% over 8 min, 70–90% over 0.6 min and 90% over 5.4 min. Mass spectrometer settings for MS analyses were a capillary voltage of 0.5 kV and a cone voltage of 40 V. The mass spectra were recorded over a scan range of 100−2000 Da. MS data were acquired using a data-dependent acquisition method (DDA) for which MS/MS data were acquired using collision energies based on mass and charge state of the candidate ions. For calibration, an external lock mass was used with a separate reference spray (LockSpray) using a solution of leucin enkephalin eluted at a flow rate of 5 μL/min. The calibration was based on the detection in MS of ions m/z 278.1141 and 556.2771 at a collision energy of 23 eV. LC/MS and LC-ESI/MS/MS data analyses were performed using MassLynx version 4.1 (Waters) software supplied by the manufacturer. The resulting MS/MS spectra data were analysed to provide *de novo* sequencing information using PEAKS^®^ studio version 5.2 software (Bioinformatics Solutions Inc.) with the following settings: chymotrypsin or Lys-C enzyme and carbamidomethylation (C) as fixed modifications, and C-terminal amidation as variable; mass accuracy on fragment ions at 0.1 Da; mass accuracy for the precursor mass at 10 ppm. The identification of peptides was further manually validated using MS/MS spectra.

### Edman degradation

Purified peptides were subjected to Edman degradation on a gas-phase sequencer model ABI 492 (Applied Biosystems, CA, USA). The phenylthiohydantoin (PTH) amino acid derivatives generated at each sequence cycle were identified and quantified on-line with an Applied Biosystems Model 140C HPLC system using the Applied Biosystems Model 610 A data analysis system for protein sequencing. The PTH-amino acid standard kit was used and reconstituted according to the manufacturer’s instructions. The procedures and reagents were used as recommended by the manufacturer.

### Chemical synthesis and purification of peptides

Fmoc amino acids were purchased from Iris Biotech GmbH (Marktredwitz, Germany). Fmoc-Gln(Trt)-Wang resin (loading 0.29 mmol/g) and Fmoc-Rink-ProTide resin (loading 0.19 mmol/g) were purchased from CEM Corporation (NC, USA). Fmoc-S-acetamidomethyle-L-cysteine (Fmoc-L-Cys(Acm)-OH) was purchased from Chem-Impex International. ACN was from Merck (Bayswater, Australia). *N*,*N*-dimethylformamide (DMF), TFA and diethyl ether were from Chem-Supply (Gillman, Australia). All solvents were of the highest available purity and used without further purification. All other reagents and solvents were obtained from Sigma-Aldrich (Sydney, NSW, Australia) in the highest available purity. Analytical RP-HPLC was performed on a Shimadzu LC-20AT system with a Kromasil Classic LC-MS C_18_ column (100 Å, 3.5 μm, 150 mm x 2.1 mm). Preparative HPLC was performed on a Vydac Protein and Peptide C_18_ preparative column and crude and fractions analysed using RP-HPLC (Shimadzu LC-20AT system) and ESI-MS. Mass analysis of the final products were performed on an API Q-star Pulsar Q-TOF mass spectrometer (PE SCIEX, Canada) with a Series 1100 solvent delivery system equipped with an auto-injector (Agilent Technologies Inc., Palo Alto, CA) and a Kromasil Classic LC-MS C_18_ column (100 Å, 3.5 μm, 150 mm x 2.1 mm). Data acquisition and processing were carried out using Analyst QS software v1.1 (PE SCIEX, Canada).

The Cys(Acm) protected A- and B-chain peptides used to make synthetic Δ-PSDTX-Pp1a and the parallel A-A homodimer were assembled using Fmoc-SPPS on a CEM Liberty Prime microwave peptide synthesiser. The A-chain with its C-terminal amide was synthesized on an Fmoc-Rink-ProTide resin (scale 0.1 mmol). The B-chain with its C-terminal acid was synthesized on a preloaded Fmoc-Gln(Trt)-Wang resin (scale 0.1 mmol). Directed disulfide-bond formation was achieved using Acm-protected cysteine building blocks, where Cys1 on the A-chain and Cys2 on the B-chain were protected with Acm. Prior to the first amino acid coupling, the Fmoc group was removed *via* treatment with 25% pyrrolidine/DMF at 105 °C for 40 s. The resin was washed with DMF (2x 4 mL). Amino acid activation and couplings were carried out in DMF using Fmoc-amino acid/carbodiimide (DIC)/Oxyma (5:10:5 equivalents of resin loading) at 105 °C for 1 min. The cycle of deprotection, washing and coupling was repeated until the full-length peptide was obtained after which the resin was washed with DCM (3x) and drained. The peptide was cleaved from the resin and the side chain protecting groups (except for Acm) were removed by treatment with 15 mL TFA: water: ethandithiol: triisopropylsilane (92.5:2.5:2.5:2.5) for 40 min at 42 °C after which the cleavage solution was drained. The crude peptide was precipitated with 30 mL cold diethyl ether, centrifuged and the supernatant discarded (repeated 3x) and redissolved in 50% ACN / 0.043% TFA in water and lyophilised. The crude peptide was purified by preparative RP-HPLC and identity confirmed using analytical RP-HPLC and ESI-Q/MS. In total, 30.1 mg A chain and 61.8 mg B chain was obtained (>90% purity) with a yield of 15% and 62%, respectively.

The Cys(Acm)-protected B chain used to make the parallel B-B homodimer was synthesized using standard Fmoc-SPPS on a Symphony (Protein Technologies Inc.) automated synthesizer using Fmoc-Gln(Trt)-Wang resin (scale 0.1 mmol). Prior to first amino acid coupling, the Fmoc group was removed *via* treatment with 30% piperidine in DMF (1 x 1.5 min and 1 x 4 min) and subsequently washed with DMF. Amino acids were coupled with 0.2 M (O-(6-Chlorobenzotriazol-1-yl)-N,N,N’,N’-tetramethyluronium hexafluorophosphate (HCTU) in DMF and N,N-diisopropylethylamine (DIEA) using 5-fold excess relative to resin loading (1 x 5 min then 1 x 10 min). The cycle of deprotection, washing and coupling was repeated until the full-length peptide was obtained after which the resin was washed with DCM (3x) and drained. Peptides were cleaved off the resin and side chain protecting groups were removed by treatment with TFA: water: ethandithiol: triisopropylsilane (90:5:2.5:2.5) for 2 h at 25° C. Following removal of most of the cleavage solvent under a stream of nitrogen, the crude peptide was precipitated with 30 mL cold diethyl ether, then the precipitate was washed with cold diethyl ether, redissolved in 50% ACN / 0.043% TFA in water, and lyophilised. The crude peptide was purified by preparative RP-HPLC and identity confirmed using analytical RP-HPLC and ESI-Q/MS. In total, 18.7 mg B chain was obtained (>90% purity) with a yield of 9%.

### Oxidative folding

Cys(Acm)-protected A-chain and B-chain peptide stock were diluted to 0.5 mM in a NH_4_HCO_3_ 0.1 M solution (pH ~ 8.0) in a 1:1.5 ratio (A-chain: B-chain) and stirred at 25 °C for 48 h to form the first disulfide bond. The reaction was monitored by analytical RP-HPLC and disulfide-bond formation confirmed by ESI-MS. Three major products were obtained: the A-B chain heterodimer, A-A homodimer and B-B homodimer, all with the Cys(Acm) protecting groups intact. The oxidative mixture was purified using preparative RP-HPLC and lyophilized.

The A-B chain heterodimer with the Cys(Acm) intact was diluted to 0.1 mM in 80% aqueous MeOH 0.1 M HCl solution to form the second disulfide bond. Fifty equivalents of I_2_ dissolved in 100% MeOH was added and the solution was stirred at 25°C for 10–20 min until the reaction was complete. The B-B homodimer second disulfide bond was formed using the same method as the A-B heterodimer. The A-A homodimer with the Cys(Acm) intact was diluted to 0.1 mM in 50% aqueous AcOH 2 M GnHCl solution then 50 equivalents of I_2_ dissolved in 50% aqueous AcOH was added and the solution was stirred at 25°C until the reaction was complete. The reactions were monitored by LC-MS. Once completed, 1 M ascorbic acid in water was added until the solution was clear. The solution was diluted 10 times with 0.043% TFA in water and the final peptide product was isolated by RP-HPLC, identity confirmed by MS and lyophilised. In total, 9.4 mg synthetic Δ-PSDTX-Pp1a (100% purity), 1.8 mg parallel A-A homodimer (97% purity), and 0.9 mg parallel B-B homodimer (92% purity) were obtained (**Table S7**).

Reduced linear unprotected A- and B-chain peptides were synthesized, purified and analysed using the same method as described for the Cys(Acm)-protected A- and B-chains. 3 mg linear A-chain and 5.5 mg linear B-chain were obtained (>95% purity) with yields of 1% and 6%, respectively (**Table S7**).

### RP-HPLC co-elution study of synthetic Δ-PSDTX-Pp1a and native Δ-PSDTX-Pp1a

RP-HPLC was performed using a Shimadzu LC-20AT system and a Kromasil Classic LC-MS C_18_ column (100 Å, 3.5 μm, 150 mm x 2.1 mm). Solvents for RP-HPLC consisted of 0.043% TFA/ water (solvent A) and 0.043% TFA/ 90% ACN/ water (solvent B). ACN was from Merck (Bayswater, Australia). TFA was from Chem-Supply (Gillman, Australia).

The crude venom, native Δ-PSDTX-Pp1a, and synthetic Δ-PSDTX-Pp1a were run on analytical RP-HPLC using a gradient of 20–70% solvent B over 25 min, at a flow rate of 0.2 mL/min. 10 μL 1 mg/mL synthetic Δ-PSDTX-Pp1a in ~15% ACN/0.043% TFA in water was injected. 40 μL crude venom in ~15% ACN/ 0.043 % TFA in water was injected. 80 μL native Δ-PSDTX-Pp1a in ~15% ACN/ 0.043% TFA in water was injected.

### Molecular modelling

Molecular models were produced using the *de novo* PEP-fold method [15] through the dedicated webserver (v.3.1 available at http://mobyle.rpbs.univ-paris-diderot.fr/cgi-bin/portal.py#forms::PEP-FOLD3), given the peptide sequences identified by MS. All structural models were submitted to molecular dynamics (MD) simulations using the Amber forcefield as implemented in Yasara [15]. Peptides were simulated in a neutralized explicit water solvent box, under periodic boundary conditions and at a constant temperature of 25°C. MD trajectories of 40 ns were collected at 2 ps intervals for both molecular systems and the production period used for analysis was set after the MD simulation reach an equilibrated state (stable root mean square deviation). Clustering analysis upon trajectories provided the most representative structure later considered as the final structural models. Electrostatic potentials were computed by using the APBS program [62] and hydrophobic potentials were provided by the Platinum webserver (http://model.nmr.ru/platinum/,[63]).

### Circular dichroism analysis of Δ-PSDTX-Pp1a

Stock peptide solutions were prepared in 50% ACN/water at 1 mM concentration. Peptide concentrations for electronic circular dichroism (ECD) analysis were 50 μM in PBS buffer (pH 7.4). ECD spectra were obtained on a Jasco J-810 spectropolarimeter (Easton, MD, USA). All experiments were carried out using a 0.1 cm quartz cell with 250 μL sample volume at 25°C. Spectra were acquired in in far UV region (185–260 nm) using 20 nm/min scan speed, 1 nm bandwidth, and 0.5 nm data pitch with 5 scans averaged for each sample. Blank subtraction was performed using Spectra Management Software followed by curve smoothing using the binomial method. Data were processed and displayed using Prism 7 (GraphPad, La Jolla USA).

### Proteolytic and thermal stability assays

Samples were analysed *via* LC-MS using an API Q-Star Pulsar mass spectrometer (SCIEX, Ontario, Canada) with a Series 1100 solvent delivery system equipped with an auto-injector (Agilent Technologies Inc., Palo Alto, CA) using a Kromasil Classic LC-MS C_18_ column (100 Å, 3.5 μm, 150 mm x 2.1 mm). Data acquisition and processing were carried out using Analyst software v1.1 (SCIEX, Canada). Solvents for RP-HPLC consisted of 0.043% TFA/ water (Solvent A) and 0.043% TFA/ 90% ACN/ water (Solvent B).

### Proteolytic stability

70 μL of reaction buffer (0.2 M sodium phosphate, 1 mM CaCl_2_ pH 7.5) was preheated to 37 °C for 30 min. 20 μL peptide stock solution (0.5 mM peptide in solvent A) and 10 μL proteinase K stock solution (5 μM in reaction buffer) were added to the reaction buffer and the mixture was incubated at 37 °C for 24 h. Final peptide concentration was 100 μM and final proteinase K concentration was 0.5 μM (1:200 proteinase K:peptide molar ratio). For the negative control, 10 μL reaction buffer was added instead of proteinase K. 10 μL aliquots were taken at 1 min, 20 min, 1 h, 2 h, 5 h, 8 h, and 24 h and quenched with 35 μL ice-cold extraction buffer (50% ACN, 0.1 M NaCl, 1% TFA) followed by centrifugation at 17,000 g for 10 min. The negative control was measured at 0 h and 24 h. Samples were stored at −30°C until they were analysed *via* LC-MS using a gradient of 20–90% solvent B over 25 min, at a flow rate of 0.2 mL/min flow. N = 2 for synthetic Δ-PSDTX-Pp1a A chain, B chain, and A-A homodimer, N = 1 for B-B homodimer (due to limited amounts of peptide). Each experiment was run in duplicate with a negative control. Peptide values were quantified relative to the negative control at 0 h and data analysis was performed using a non-linear fit one-phase decay model in Prism Version 8 (GraphPad, La Jolla USA).

### Thermal stability

Δ-PSDTX-Pp1a was incubated for 72 h at 37°C, 60°C, and 90°C. 48 μL 0.05% TFA in water was preheated to 37°C, 60°C, or 90°C for 30 min. Once heated, 0.5 mM Δ-PSDTX-Pp1a was diluted in the heated buffer to a final concentration of 100 μM and a total reaction volume of 60 μL. Time points were taken at 0 h, 24 h, 48 h, and 72 h and at each timepoint 10 μL reaction solution was diluted in 35 μL 0.05% TFA in water and stored at −30°C until analysis *via* LC-MS using a gradient of 20–90% solvent B over 18 min, at a flow rate of 0.2 mL/min. Peptide values were quantified by ion extraction (ion range 1100.6–1101.6 Da) using Analyst QS 1.1 software quantification wizard. N = 3 for each peptide. Experiments were run in triplicate and samples were quantified relative to t=0 h. Data analysis was performed using a non-linear fit one-phase decay model in Prism Version 8 (GraphPad, La Jolla USA). Y0 and Plateau were constrained to 100 and 0, respectively.

### *In vivo* insecticidal assays

Insecticidal activity was evaluated injection of peptides into the ventro-lateral thoracic region of sheep blowflies (*Lucilia cuprina*; mass 22.4–31.7 mg) as previously described [60]. A 1.0 mL Terumo Insulin syringe (B-D Ultra-Fine, Terumo Medical Corporation, Maryland, USA) with a fixed 29 G needle fitted to an Arnold hand micro-applicator (Burkard Manufacturing Co. Ltd., England) was used for injection with a maximum volume of 2 μL per fly. All flies were individually housed in 2 mL tubes and paralytic activity and lethality were determined at 0.5, 1 and 24 h post-injection. A total of three tests were performed with 4–6 doses per peptide (n = 10 flies per dose) and the corresponding controls (MilliQ water; n = 10–40 per peptide). PD_50_ and LD_50_ values were calculated as previously described [64]. Two-way ANOVA with Tukey’s multiple post hoc test was performed in Prism 8 for statistical comparison of the insecticidal activity of different peptides. PD_50_ and LD_50_ values for the A- and B-chain monomers and homodimers were compared to the values of the heterodimeric Δ-PSDTX-Pp1a (* p<0.01; ** p<0.0001).

## Acknowledgments

This work was supported by Investissement d’Avenir of the Agence National de la Recherche (CEBA: ANR-10-LABX-25-01). We are grateful to the Virology Unit and the Medical Entomology Unit of the Pasteur Institute of French Guiana for kindly providing *Aedes albopictus* cells and *Aedes aegypti* eggs and we like to thank Geoff Brown (Department of Agriculture and Fisheries, Queensland, Australia) for the supply of blowflies. M.M. was supported by the European Research Council under the European Union’s Horizon 2020 research and innovation program (714366), by the Australian Research Council (DP190101667), and by the Vienna Science and Technology Fund (WWTF; LS18-053). G.F.K. was supported by a Principal Research Fellowship (APP1136889) from the Australian National Health & Medical Research Council.

## Author contributions

A.T., A.D., M.M. and C.D. conceived the study. A.T. collected and dissected ants. I.B. performed the cytotoxic and fluorometry bioassays. The peptide purification and MALDI-TOF MS/MS analysis was made by M.T., L.J. and data were analysed by L.J., M.D.W, R.B. and AT. Insecticidal assays were conducted by V. H. H.M. synthesized, purified synthetic peptides and performed the stability tests under the supervision of M.M. ECD spectra were generated by N.B.E. Structural modelling was performed by O.D and A.T. optimized the visualization. All authors discussed the results and contributed to the final manuscript.

