## Supplementary Information for "Heterodimeric insecticidal peptide provides new insights into the molecular and functional diversity of ant venoms"

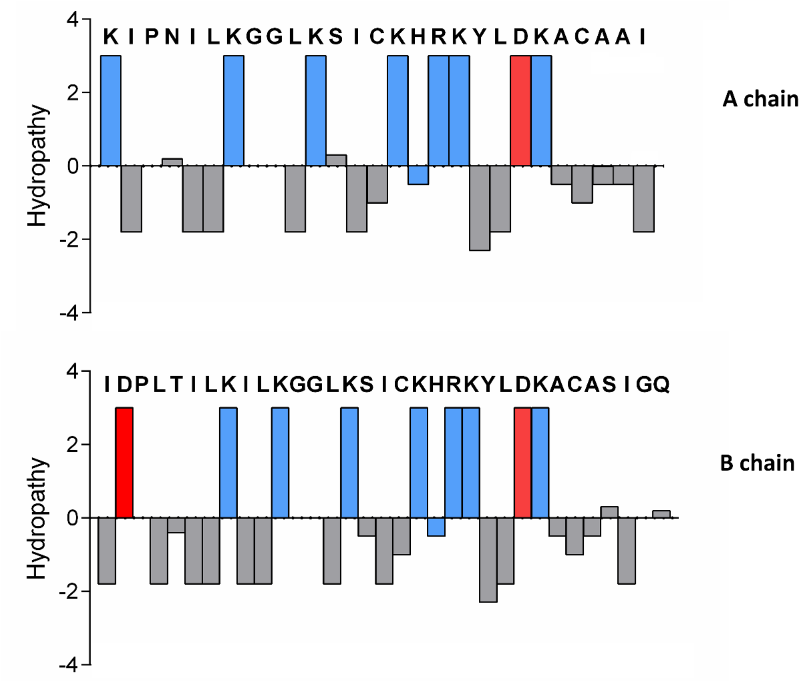

**Figure S1. Physicochemical properties of Δ-PSDTX-Pp1a chains.** Δ-PSDTX-Pp1a is comprised of two amphiphilic and polycationic chains. Hydropathy plot for Δ-PSDTX-Pp1a chains by using hydrophobicity values for each amino acid (1). Hydrophilicity is shown as positive and hydrophobicity as negative relative to the central mid-line. Positively charged residues are indicated in blue while negatively charged amino acids are in red.

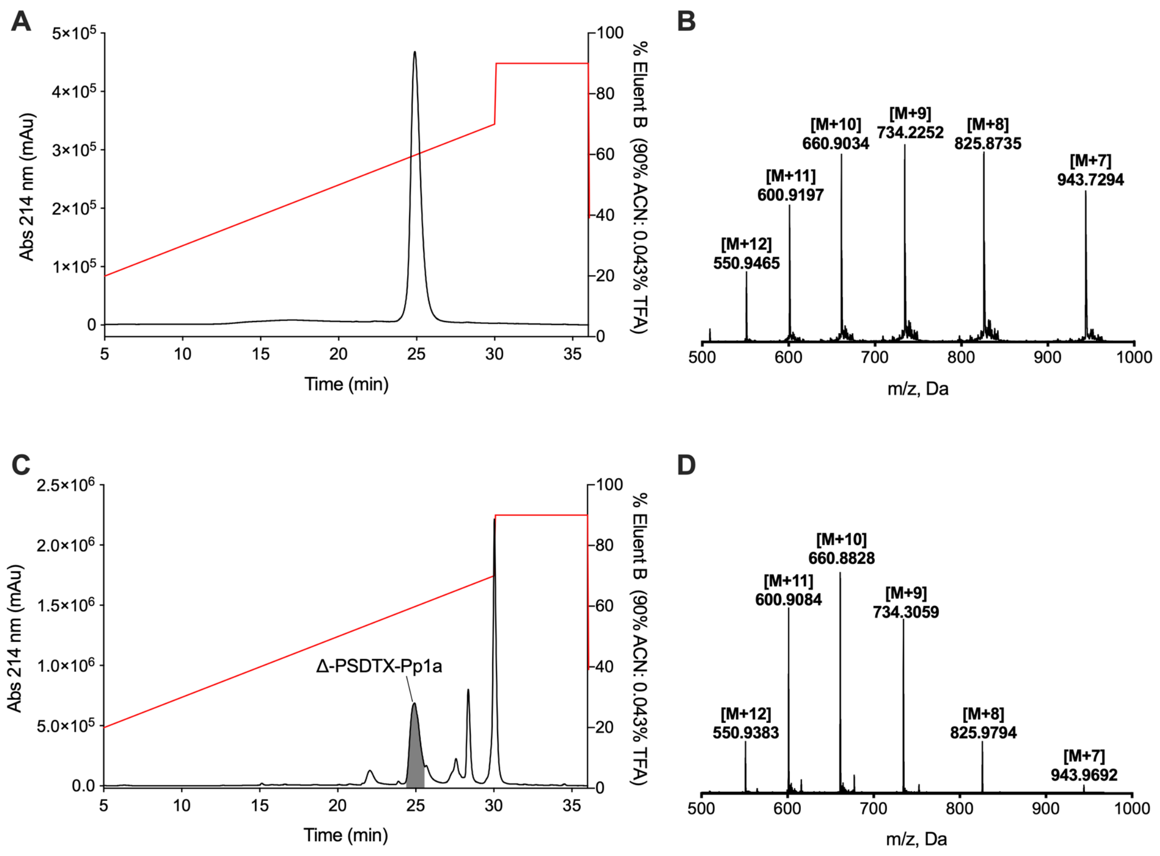

**Figure S2. HPLC-MS profiles of synthetic Δ-PSDTX-Pp1a and crude ant venom.** Analytical RP-HPLC chromatograms of (**A**) synthetic Δ-PSDTX-Pp1a and (**C**) crude ant venom (native Δ-PSDTX-Pp1a is highlighted) obtained with a Shimadzu LC-20AT system using a Kromasil Classic LC-MS $C_{18}$ column (100 Å, 3.5 µm, 150 mm x 2.1 mm) with a gradient of 20–70% solvent B over 25 min, with a flow rate of 0.2 mL/min. High-resolution mass spectra of (**B**) synthetic Δ-PSDTX-Pp1a (calculated monoisotopic mass 6598.78 Da; measured monoisotopic mass 6598.8 Da) and (**D**) native Δ-PSDTX-Pp1a (calculated monoisotopic mass 6598.8 Da, measured monoisotopic mass 6598.6 Da) obtained using a Q-Star Elite ESI Q-TOF mass spectrometer (PE SCIEX, Canada).

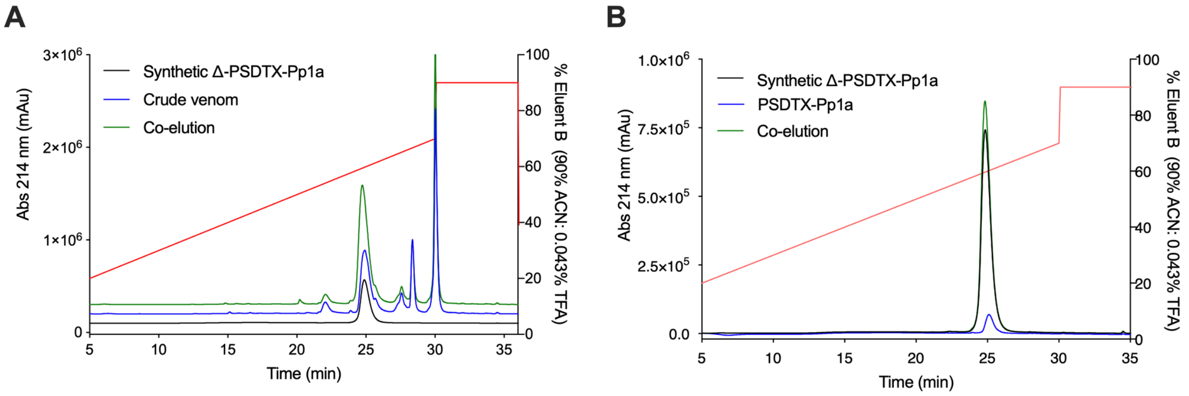

**Figure S3. RP-HPLC co-elution study of synthetic Δ-PSDTX-Pp1a and native Δ-PSDTX-Pp1a**. (**A**) Overlaid RP-HPLC chromatograms of synthetic Δ-PSDTX-Pp1a, crude ant venom, and a mixture of both. (**B**) Overlaid RP-HPLC chromatograms of synthetic and native Δ-PSDTX-Pp1a purified from venom, and a mixture of both.

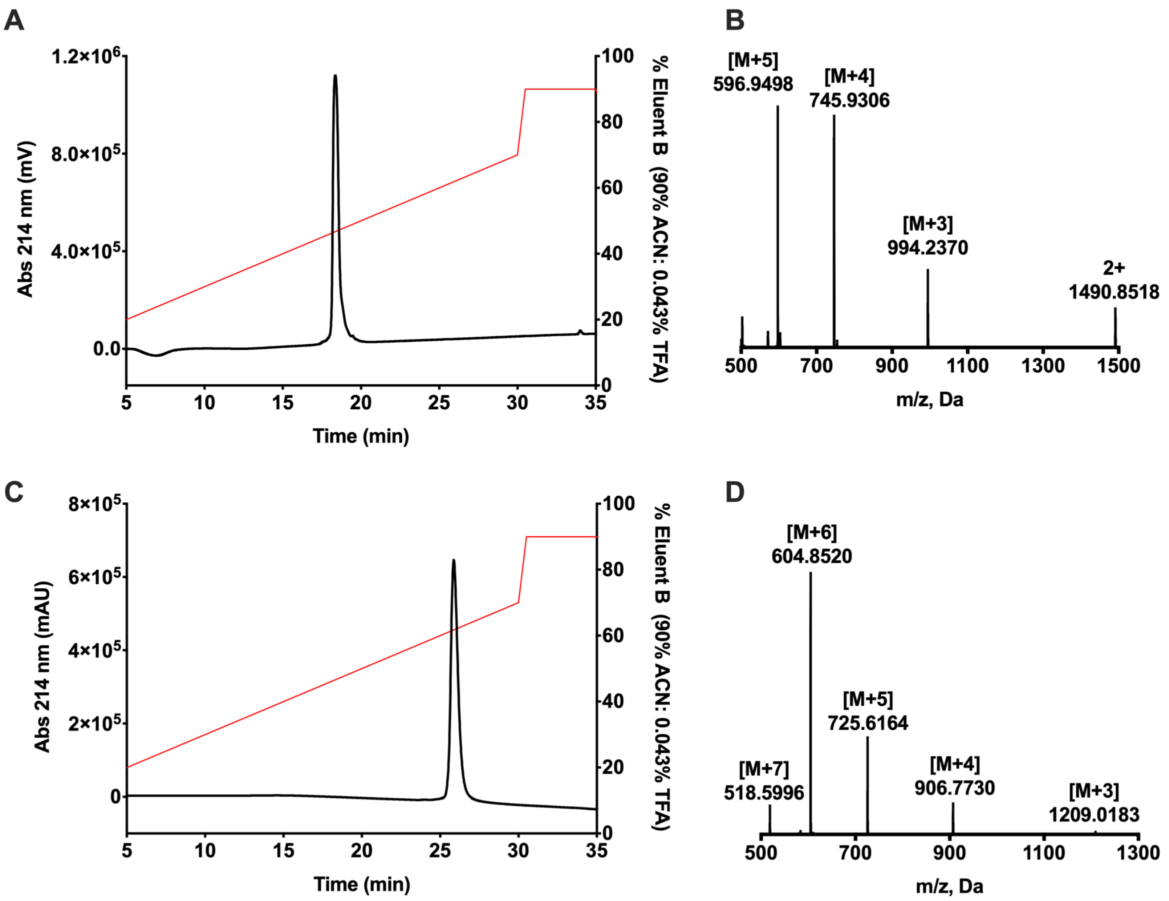

**Figure S4: HPLC-MS profiles of reduced linear A-chain and B-chain monomer.** Analytical RP-HPLC chromatograms of **(A)** A-chain monomer and **(C)** B-chain monomer obtained with a Shimadzu LC-20AT system using a Kromasil Classic LC-MS C18 column (100 Å, 3.5 μm, 150 mm x 2.1 mm) with a gradient of 20–70% solvent B over 25 min, using a flow rate of 0.2 mL/min. High-resolution MS spectra of **(B)** A-chain monomer and **(D)** B-chain monomer obtained using an API Q-Star Pulsar ESI Q-TOF mass spectrometer (PE SCIEX, Canada). Measured isotope peaks are listed in Table S4.

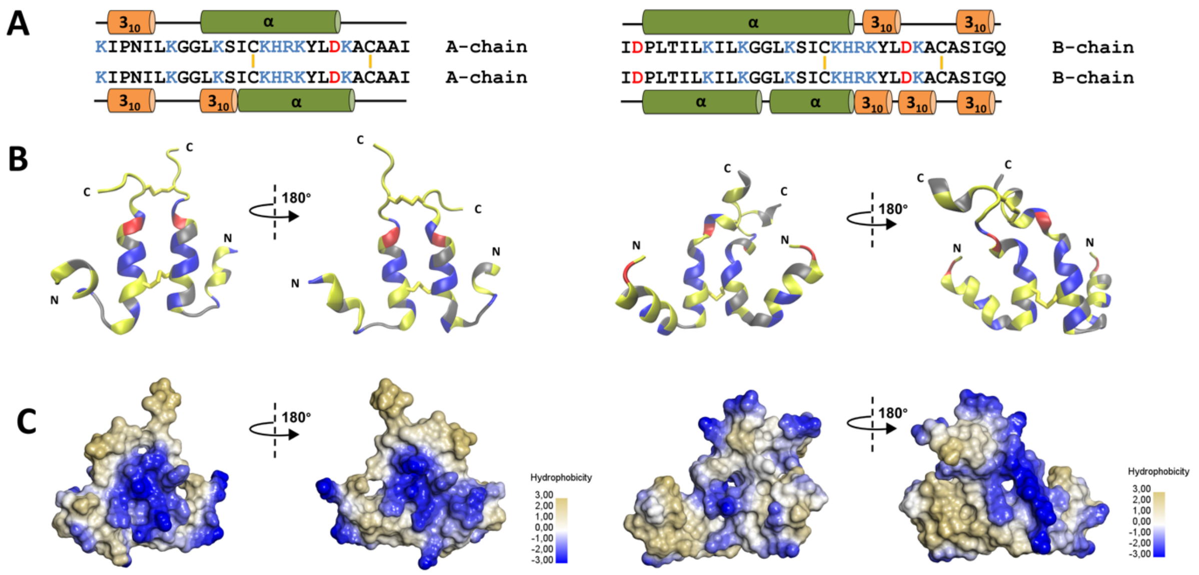

**Figure S5. Modelled structures of parallel homodimers.** Amino acid sequence of the AA **(A)** and BB **(D)** homodimer with cationic and anionic residues in blue and red, respectively. Schematic representation of the secondary structure of each homodimer is shown above and below the chain sequences. Most representative geometry for AA **(B)** and homodimer BB **(E)** homodimer after clustering of a 40 ns molecular dynamics simulation. Cationic residues are in blue, anionic residues in red, polar non-charged residues in grey, and hydrophobic residues in yellow. Molecular surface representations of the AA **(C)** and BB **(F)** homodimers**.**

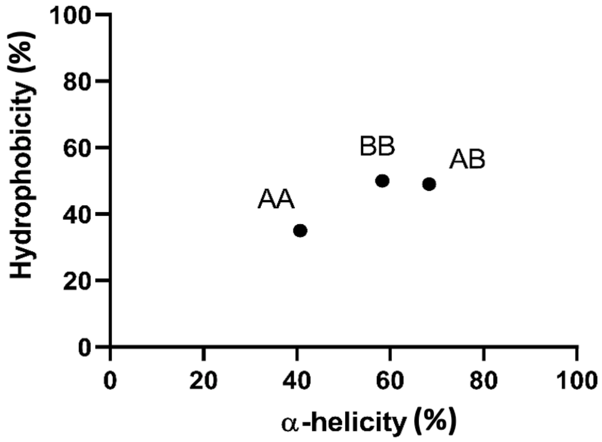

**Figure S6. Percentage of hydrophobic molecular surface plotted against percentage of α-helicity of the parallel homodimers AA, BB and the antiparallel heterodimer AB (Δ-PSDTX-Pp1a)**.

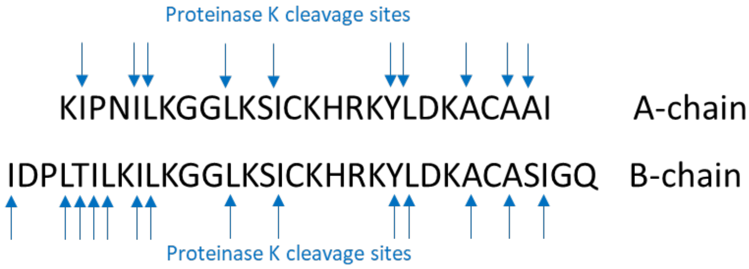

**Figure S7. Predicted proteinase K cleavage sites for Δ-PSDTX-Pp1a chains** determined using the ExPASy PeptideCutter tool (https://web.expasy.org/peptide_cutter/).

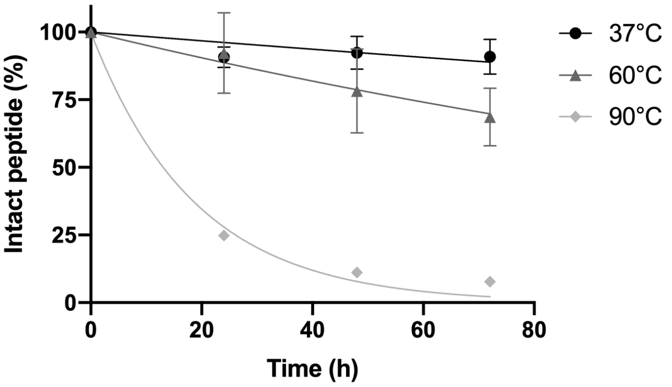

**Figure S8. Thermal stability of Δ-PSDTX-Pp1a at 37°C, 60°C and 90°C**. Δ-PSDTX-Pp1a was incubated at 100 µM and time points taken every 24 hours. Samples were analyzed using LC-MS and quantified relative to t = 0 h. Each experiment was run in triplicate. Data were fit with a non-linear fit one-phase decay model in Prism Version 8. Error bars depict the standard error of the mean (SEM). Note: error bars for the 90°C values are smaller than the symbol.

**Table S1. Monoisotopic masses of the major RP-HPLC peaks of *P. penetrator* venom and correspondence with peptides identified in a previous study (2)**

| LC-MS (this study) | | LC-MS (2) | |
|---|---|---|---|
| **Peak** | **Monoisotopic mass (Da)** | **Monoisotopic mass (Da)** | **S-S features** |
| 4 | 720.4 | – | |
| 4 | 649.4 | – | |
| 5 | 5955.4 | 5956.2 | 2 S-S (homodimer) |
| 5 | 1587.0 | – | |
| 5 | 1324.8 | – | |
| 6 | 6598.8 | 6600.9 | 2 S-S (heterodimer) |
| 6 | 1910.2 | 1911.1 | no S-S (linear) |
| 6 | 1981.3 | 1982.1 | no S-S (linear) |
| 6 | 2052.3 | – | |
| 7 | 7242.1 | 7244.3 | 2 S-S (homodimer) |
| 7 | 6598.8 | 6600.9 | 2 S-S (heterodimer) |
| 8 | 3915.4 | 3915.9 | no S-S (linear) |
| 8 | 2536.6 | – | |
| 8 | 3662.4 | – | |
| 9 | 2683.4 | 2684.3 | no S-S (linear) |
| 9 | 5254.9 | 5255.6 | no S-S (linear) |

**Table S2. Concentrations of *P. penetrator* venom extract and fractions that inhibited 50% and 99% of *A. albopictus* cell growth after 24 hours ($IC_{50}$ and $IC_{99}$).** 95% confidence interval was calculated with a logistic regression by Probit analysis on the basis of percentage of growth inhibition (Wald chi-square, Likehood Ratio chi-square and associated p-values of the regression tests are given).

| | $IC_{50}$ (µg/mL) | 95% confidence intervals | $IC_{99}$ (µg/mL) | 95% confidence interval | $X^2$ Wald (p-value) | Khi² LR (p-value) |
|---|---|---|---|---|---|---|
| Cypermethrin | 150 | 135.8–165.8 | 362.6 | 329.5–405.8 | 168.93 (<0,0001) | 250.31 (<0,0001) |
| Venom extract | 2.0 | 0.9–3.1 | 21.8 | 18.8–25.9 | 140.18 (<0.0001) | 407.62 (<0.0001) |
| Fraction* 4 | >70 | – | >70 | – | 1.33 (0.311) | 3.66 (0.099) |
| Fraction 5 | 43.2 | 7.6–46.8 | 64.3 | 57.4–141.5 | 5.42 (0.020) | 459.22 (<0.0001) |
| Fraction 6 | 3.2 | 3.0–3.4 | 5.9 | 5.5–6.4 | 439.79 (<0.0001) | 958.17 (<0.0001) |
| Fraction 7 | 28.9 | 25.3–33.0 | 60.5 | 54.5–68.0 | 238.30 (<0.0001) | 537.41 (<0.0001) |
| Fraction 8 | 29.5 | 19.0–42.5 | 58.4 | 45.2–74.4 | 1295.90 (<0.0001) | 505.15 (<0.0001) |
| Fraction 9 | 30.5 | 20.1–44.0 | 57.2 | 43.8–74.2 | 434.52 (<0.0001) | 543.65 (<0.0001) |

**Fractions 4-9 correspond to those identified in the HPLC chromatogram in Figure S1*

**Table S3. Peptide fragments observed in LC-MS/MS analysis of the enzymatic digest of carbamidomethyl derivates of each monomer of Δ-PSDTX-Pp1a**. Bold "C" denotes carbamidomethyl derivatization of cysteine residues.

| | Fragments | *Observed m/z* | Relative error ppm* |
|---|---|---|---|
| **Δ-PSDTX-Pp1a A-chain** | KIPNILKGGLKSICKHRKYLDKACAAI | | |
| | KHRKY | 366.2187 | -1.4 |
| | KGGLKSI**C**KH | 376.5498 | -0.5 |
| | KGGLKSI**C** | 431.7446 | 0.5 |
| | SI**C**K | 254.1338 | 1.5 |
| | YLDK | 269.6478 | 2.3 |
| | KIPNIL | 413.2811 | -0.8 |
| **Δ-PSDTX-Pp1a B-chain** | IDPLTILKILKGGLKSICKHRKYLDKACASIGQ | | |
| | IDPLTILK | 456.7920 | 0.2 |
| | LDKA**C**ASIGQ | 531.7703 | 7.9 |
| | A**C**ASIGQ | 353.6624 | -1.9 |
| | ILKGGLK | 364.7551 | 0.0 |
| | SI**C**K | 254.1342 | 3.1 |

*Related to the calculated mass*

**Table S4. High-resolution mass spectrometry measured isotope peaks for synthetic peptides**.

| m/z | Δ-PSDTX-Pp1a | A-A homodimer | B-B homodimer | A-chain monomer | B-chain monomer |
|---|---|---|---|---|---|
| 2+ | Not detected | Not detected | Not detected | 1490.8518 | Not detected |
| 3+ | Not detected | Not detected | Not detected | 994.2370 | 1209.0183 |
| 4+ | Not detected | Not detected | Not detected | 745.9306 | 906.7730 |
| 5+ | Not detected | Not detected | Not detected | 596.9498 | 725.6164 |
| 6+ | 1100.8538 | 994.237 | 1208.1765 | Not detected | 604.8520 |
| 7+ | 943.7294 | 852.3481 | 1035.5677 | Not detected | 518.5996 |
| 8+ | 825.8735 | 745.9306 | 906.2473 | Not detected | Not detected |
| 9+ | 734.3353 | 663.0466 | 805.6658 | Not detected | Not detected |
| 10+ | 660.9197 | 596.9406 | 725.2045 | Not detected | Not detected |
| 11+ | 600.9197 | 542.9562 | 659.3701 | Not detected | Not detected |
| 12+ | 550.9465 | Not detected | 604.6752 | Not detected | Not detected |

**Table S5. Insecticidal effects in blowflies.** All experiments were performed using three repeats per peptide as detailed in the Materials and Methods section.

| | **$PD_{50}$-0.5h (nmol/g)** | | **$PD_{50}$-1h (nmol/g)** | | **$PD_{50}$-24h (nmol/g)** | | **$LD_{50}$-24h (nmol/g)** | | **Comment** |
|---|---|---|---|---|---|---|---|---|---|
| | av. | SEM | av. | SEM | av. | SEM | av. | SEM | |
| **A-B heterodimer** | 2.5 | 0.6 | 2.4 | 0.6 | 2.4 | 0.6 | 3.0 | 0.5 | Caused contractile paralysis on the needle within seconds of injection, then lethal |
| **B-B homodimer** | 3.7 | 0.5 | 3.2 | 0.2 | 4.3 | 0.0 | 4.6 | 0.2 | Caused contractile paralysis on the needle within seconds of injection, then lethal |
| **A-A homodimer** | 12.7 | 1.2 | 10.7 | 0.1 | 14.3 | 0.5 | 16.1 | 0.4 | Caused contractile paralysis on the needle within seconds of injection, then lethal |
| **B-chain** | 21.7 | 1.1 | 15.4 | 2.1 | 29.4 | 2.3 | 35.8 | 0.1 | Caused contractile paralysis first, then lethal |
| **A-chain** | 70.4 | 2.2 | 50.6 | 2.1 | 92.9 | 12.3 | 108.6 | 10.1 | Only active at very high doses |

*$PD_{50}$ = dose that causes paralysis in 50% of flies, $LD_{50}$ = dose that causes lethality in 50% of flies, av. = average, SEM = standard error of the mean.*

**Table S6. Physicochemical characteristics of dimeric ant-venom peptides.**

| Peptide | Sub Family | Structure | S-S | hydrophobic aa | Net charge | Mass | Target |
|---|---|---|---|---|---|---|---|
| Δ-PSDTX-Pp1a | PM | heterodimer | 2 | 43% | + 13 | 5598.74 Da | Unknown |
| $U_1$-PSDTX-Pt1a | PM | heterodimer | 2 | 47% | + 8 | 6940.90 Da | Unknown |
| $U_1$-PSDTX-Pt1b | PM | heterodimer | 2 | 47% | + 9 | 6864.85 Da | Unknown |
| $U_1$-PSDTX-Pt1c | PM | heterodimer | 2 | 47% | + 8 | 6968.90 Da | Unknown |
| $U_1$-PSDTX-Pt1d | PM | heterodimer | 2 | 47% | + 8 | 7025.93 Da | Unknown |
| $U_1$-PSDTX-Pt1e | PM | heterodimer | 2 | 47% | + 9 | 6892.85 Da | Unknown |
| $U_1$-PSDTX-Pt1f | PM | heterodimer | 2 | 47% | + 9 | 6949.87 Da | Unknown |
| $U_2$-PSDTX-Ta1a | PM | homodimer | 1 | 40% | + 6 | 5750.94 Da | Unknown |
| $U_2$-PSDTX-Ta1b | PM | homodimer | 3 | 44% | + 6 | 5438.60 Da | Unknown |
| $U_2$-PSDTX-Ta1c | PM | homodimer | 1 | 40% | +4 | 5608.78 Da | Unknown |
| $U_2$-PSDTX-Ta1a/c | PM | heterodimer | 1 | 40% | +5 | 5680.86 Da | Unknown |
| M-MIITX-Mp2a | M | heterodimer | 2 | 50% | + 13 | 5603.28 Da | Membrane cell |
| $U_1$-MIITX-Mp2b | M | heterodimer | 2 | 49% | + 13 | 5660.30 Da | Unknown |
| $U_1$-MIITX-Mp3a | M | homodimer | 2 | 42% | + 14 | 8192.60 Da | Unknown |
| $U_1$-MIITX-Mp4a | M | homodimer | 1 | 49% | + 6 | 8540.82 Da | Unknown |
| $U_1$-MIITX-Mg2a | M | homodimer | 1 | 51% | +12 | 8482.78 Da | Unknown |
| $U_1$-MIITX-Mg4b | M | homodimer | 5 | 43% | +14 | 9220.85 Da | Unknown |
| ω/M-ECTX-Et1a | E | heterodimer | 3 | 44% | + 9 | 7923.17 Da | Membrane cell |
| $U_1$-ECTX-Et1b | E | heterodimer | 3 | 51% | + 13 | 8208.55 Da | Unknown |
| $U_1$-ECTX-Eb1a | E | heterodimer | 1 | 53% | +17 | 9011.20 Da | Unknown |
| $U_1$-ECTX-Eb1b | E | homodimer | 1 | 53% | +18 | 9419.45 Da | Unknown |
| $U_1$-PONTX-Om4a | P | homodimer | 1 | 48% | +10 | 6331.63 Da | Unknown |

*PM: Pseudomyrmecinae, M: Myrmeciinae, E: Ectatomminae, P: Ponerinae, aa: amino acids*

**Table S7. Molecular mass and measured ions for dimeric ant-venom peptides.**

| Peptide | Molecular weight | Observed mass ion | Calculated mass | Observed mass |
|---|---|---|---|---|
| Synthetic Δ-PSDTX-Pp1a | 6598.78 | 7+ | 943.691 | 943.765 |
| A chain linear homodimer | 5955.44 | 6+ | 993.588 | 994.237 |
| B chain linear homodimer | 7242.13 | 6+ | 1208.035 | 1208.128 |
| A chain monomer | 2979.74 | 3+ | 994.255 | 994.231 |
| B chain monomer | 3623.08 | 4+ | 906.778 | 906.782 |
